# Intrinsic higher potency of basal urothelial cells to intermediate and umbrella cells as the cell of origin for bladder cancer

**DOI:** 10.1101/2025.06.28.662140

**Authors:** Chuan Yu, Nicholas Chu, Alexandra Aguirre, Jonathan Green, Qing Xie, Beatrice Knudsen, Zhu A. Wang

## Abstract

Transcriptome profiling of bladder cancer has revealed distinct basal-like and luminal-like molecular subtypes, which may be correlated with pathological subtypes of different patient outcomes. However, whether these molecular subtypes originate from the corresponding cell types in the normal urothelium and whether different cells of origin influence bladder cancer progression remain unclear. Here, we conducted cell-type-specific lineage tracing in CRISPR/Cas9-induced mouse bladder cancer models of *Pten* and *Trp53* targeting. We show that although basal, intermediate, and superficial umbrella cells can all serve as the cell of origin for bladder cancer, transformed umbrella cells were gradually displaced by tumor cells from inner layers, particularly transformed basal cells, which had highest stemness. Histological and single cell RNA-sequencing data comparing basal- and intermediate-cell-induced bladder tumors revealed that basal-induced tumors displayed higher heterogeneity, and contained unique cell clusters including Krt14+Ki67+ highly proliferative basal cells, squamous cell carcinoma, and transitioning cells towards the Gata3+ luminal subtype. Trajectory analysis confirmed the cell lineage differentiation hierarchy uncovered in lineage tracing. Moreover, human bladder cancer molecular subtype signatures were highly enriched in mouse tumor cell clusters of the corresponding cell of origin, and a gene signature derived from the unique basal-induced clusters is predictive of worse patient outcome. Overall, our results support that the basal and luminal molecular subtypes of bladder cancer have the corresponding cells of origin as their basis, and that urothelial basal cells are intrinsically more competitive than intermediate and umbrella cells in generating aggressive bladder cancer subtypes.

## Introduction

Bladder cancer is a clinically heterogeneous disease, with ∼90% of cases being urothelial carcinomas originating from bladder epithelium precursor lesions. Pathologically, bladder cancer can be categorized into non-muscle invasive bladder cancer (NMIBC, ∼80% of initial cases) and muscle invasive bladder cancer (MIBC, ∼20%), which is the more aggressive subtype (Knowles and Hurst, 2015; Guo et al., 2024). Transcriptomic profiling has characterized distinct molecular subclasses in both NMIBC and MIBC, broadly defined as the basal-like and luminal-like subtypes (Cancer Genome Atlas Research, 2014; Choi et al., 2014a; Choi et al., 2014b; Hedegaard et al., 2016; Robertson et al., 2017). Concomitantly, genomic and integrated multi-omics characterization of bladder cancers also implies that basal and luminal molecular subtypes may be associated with distinct pathological subtypes and different patient outcomes (Knowles and Hurst, 2015; Hedegaard et al., 2016; Glaser et al., 2017; Hurst et al., 2017; Hurst et al., 2021; Lindskrog et al., 2021; de Jong et al., 2023; Guo et al., 2024). This raises the intriguing question as to whether the cell of origin, defined as a normal cell type capable of generating a tumor under oncogenic conditions (Visvader, 2011; Hoadley et al., 2018), is an underlying critical factor in determining these different molecular and pathological subtypes. Answer to this question is not only important for our deeper understanding of the bladder cancer progression mechanism, but also development of novel personalized therapeutic strategies.

In both human and mouse, the bladder urothelium, enveloped by the lamina propria and detrusor muscle, is a stratified epithelial tissue composed of a bottom layer of basal cells and multiple layers of luminal cells that include intermediate cells and superficial umbrella cells (Kobayashi et al., 2015; Wiessner et al., 2022) (Fig. 1A). While the normal adult bladder urothelium is extremely quiescent, stem/progenitor cells exist to function during injury-induced tissue repair (Wiessner et al., 2022). Lineage-tracing studies and recent single cell analysis of mouse bladder regeneration indicated that intermediate cells could serve as progenitors and differentiate to produce umbrella cells (Gandhi et al., 2013; Wang et al., 2018; Cheng et al., 2021). Moreover, basal cells have also been implicated to be stem cells capable of generating all urothelial cell lineages under the injury-repair condition *in vivo* as well as in organoids and assembloids (Shin et al., 2011; Papafotiou et al., 2016; Kim et al., 2020). In analyzing the cell of origin for bladder cancer, previous research has mostly relied on the chemical-induced *N*-butyl-*N*-(4-hydroxybutyl)nitrosamine (BBN) tumorigenesis mouse model, where basal cells were shown to more likely generate aggressive carcinoma in situ (CIS) and invasive bladder tumors, whereas intermediate cells to less aggressive papillary tumors (Shin et al., 2014; Van Batavia et al., 2014; Papafotiou et al., 2016; Tate et al., 2021). Despite the progress, due to the highly variable nature of chemical carcinogenesis, it remains unclear whether those results reflected intrinsic basal cell properties as the cell of origin or the different influences of BBN on the mutational landscapes of various cell types. Analyzing the cell of origin for bladder cancer under defined genetic mutational background should more closely mimic physiological cancer-initiating conditions and help distinguish these possibilities. Here, we introduce a novel mouse bladder cancer model system by integrating the CRISPR/Cas9-induced genetic targeting with cell-type-specific lineage tracing. Our histological and single cell RNA-sequencing (scRNA-seq) molecular analyses provide definitive evidence supporting that basal cells are intrinsically more potent in serving as the cell of origin for the aggressive subtypes of bladder cancer. Our results also for the first time link the basal and luminal molecular subtypes of bladder cancer to their corresponding cells of origin.

**Figure 1.**
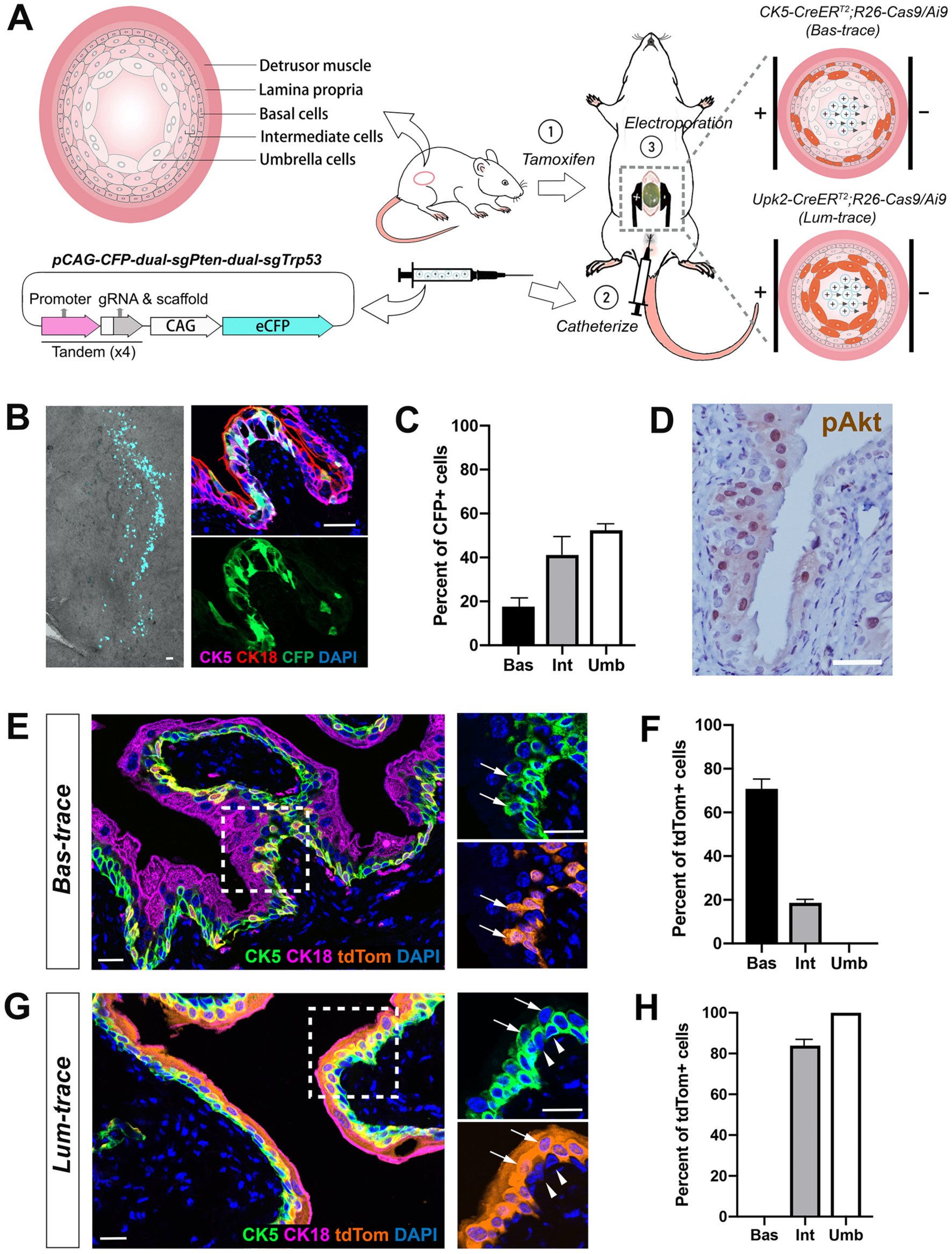
Establishment of CRISPR-based lineage-traceable bladder cancer mouse models. (**A**) Schematic diagram of labeling of basal and luminal cells in the mouse bladder urothelium before bladder tumor induction by CRISPR. (**B**) Direct visualization of CFP in the bladder after electroporation (left) and IF staining showing CFP signals in the urothelial cells but not stromal cells (right). (**C**) Quantitation of the proportions of CFP+ cells in each urothelial cell type after electroporation. N=4 animals used. (**D**) IHC showing positive pAkt signals in the urothelial cells 2 weeks post electroporation. (**E**) IF showing tdTomato labeling of basal and intermediate cells in the *Bas-trace* model. Arrows in inset point to labeled intermediate cells. (**F**) Quantitation of the proportions of tdTom+ cells in each urothelial cell type of the *Bas-trace* mice. N=4 animals used. (**G**) IF showing tdTomato labeling of umbrella and intermediate cells in the *Lum-trace* model. Arrows in inset point to labeled intermediate cells, and arrowheads to basal cells, which were unlabeled. (**H**) Quantitation of the proportions of tdTom+ cells in each urothelial cell type of the *Lum-trace* mice. N=4 animals used. Scale bars in **B**,**D**,**E**,**G** correspond to 50 µm. Erros bars in **C**,**F**,**H** correspond to one s.d.

## Results

### Establishment of CRISPR-based lineage-traceable bladder cancer mouse models

To compare the oncogenic potential of different urothelial cell types under defined genetic mutant background, we have developed novel bladder cancer mouse models that integrate CRISPR gene targeting and cell lineage tracing (Fig. 1A). Specifically, using our previously described electroporation method (Yu et al., 2018), we can deliver sgRNA-expressing plasmids specifically to the bladder urothelium of Cas9-expressing mice. Additionally, by employing the Cre-loxP-based lineage tracing strategy, we can label basal or luminal cells prior to tumor induction, and characterize the resulting tumors at both the phenotypic and molecular levels. The Trp53 and PI3K pathways are among the most frequent genomic alterations in aggressive bladder cancers (Cancer Genome Atlas Research, 2014; Hedegaard et al., 2016; Robertson et al., 2017), and *Adeno-Cre-*mediated knockout of *Pten* and *Trp53* genes in mice resulted in invasive bladder tumors (Puzio-Kuter et al., 2009). We therefore targeted these two pathways as the defined genetic mutant background for our cell-of-origin analyses. For *Pten* and *Trp53*, we designed dual sgRNAs targeting each gene and validated their efficiencies (Fig. S1A, B), so that large deletions (hence gene loss-of-function) could be generated as reported previously (Xue et al., 2014). All four sgRNAs were cloned into a single plasmid *pCAG-eCFP-dual-sgPten-dual-sgTrp53* (Fig. S1C, Table S1), which was urethral-injected and electroporated into bladder urothelial cells of R26-Cas9-EGFP (referred to as R26-Cas9) mice. Three days post electroporation, we detected CFP expression in 52.4% of umbrella cells, 41.1% of intermediate cells, and 17.6% of basal cells, but not in stromal cells of the lamina propria and muscles (Fig. 1B, C), indicating that the plasmid delivery was both efficient and specific. From the distribution of CFP-expressing cells, it was evident that the penetration rate of plasmid entering urothelial cells is correlated with the depth of the cell type relative to the lumen, with umbrella cells being the most accessible for gRNA targeting while inner basal cells the most challenging to reach (Fig. 1C). Two weeks post electroporation, we detected elevated levels of phospho-Ser473-Akt (pAkt) in the urothelium (Fig. 1D), demonstrating the efficacy of our strategy.

For tracing of cells of origin in bladder tumors, we introduced the *R26-LSL-tdTomato* (*Ai9*) reporter (Madisen et al., 2010) and the ubiquitously-expressed *R26-Cas9* line (Platt et al., 2014) to the bovine cytokeratin 5 (CK5) promoter-driven CreER^T2^ allele (Indra et al., 1999) to generate the *CK5-CreER^T2^; R26-Cas9/Ai9* mice (referred to as *Bas-trace* model) for basal cell lineage tracing. In parallel, we used the mouse uroplakin 2 promoter (Upk2)-driven CreER^T2^ knock-in allele (Shen et al., 2012) to generate the *Upk2-CreERT2; R26-Cas9/Ai9* mice (referred to as *Lum-trace* model) for luminal cell lineage tracing. To assess their specificity and efficiency, both models were tamoxifen-induced at 7-weeks of age, and analyzed one week later. For the *Bas-trace* model, 70.8% of basal cells, which are defined as CK5+CK18- and located near the basal lamina, as well as 18.6% of intermediate cells, which are located in the middle layers of the urothelium and express moderate levels of CK5 and CK18, were labeled with tdTomato. In contrast, no superficial umbrella cells, which are CK5-CK18+ and located at the apical side of the urothelium, were labeled (Fig. 1E, F). For the *Lum-trace* model, 100% of the superficial umbrella cells and 83.9% of the intermediate cells were tdTomato-labeled, whereas no basal cell was tdTomato-positive (Fig.1G, H). Although both models labeled portions of intermediate cells, the complete lack of labeling of the umbrella or basal cells in the *Bas-* or *Lum-trace* models, respectively, demonstrated high specificity of the tracing systems and enabled us to deduce the cells of origin for different tumor subtypes by comparing the two models together, as illustrated below.

### *Pten* and *Trp53* inactivation in mouse bladder urothelium induced two subtypes of tumors of distinct morphological features

Our strategy of identifying the preferred cell of origin for bladder cancer is relied upon labeling wildtype basal (plus a few intermediate) or luminal (all the umbrella plus the majority of intermediate) cells prior to tumor initiation in *Bas-trace* and *Lum-trace* mice, respectively, followed by analyzing and comparing the status of tdTomato-labeled cells between the two models after tumor initiation (Fig. 2A), similar to what we did previously for prostate cancer (Wang et al., 2014). Possible lineage-tracing outcomes are shown in Fig. 2A. For instance, in the *Lum-trace* model, if we observe that the majority of the tumor cells are tdTom-negative, it will strongly argue for the basal cells as the preferred cell of origin. We first confirmed the reliability of our CRISPR approach as all of the 36 mice that underwent electroporation developed tumors to various degrees through time (Table S2). Three months post electroporation, H&E staining revealed that the bladder epithelium contained extensive neoplastic lesions (Fig. 2B) and papillary tumors that protruded into the bladder lumen (Fig. 2C, D, Fig. S2A). The lamina propria appeared mostly normal, but in some cases, lymphocyte infiltration into the urothelial tumor could be seen (Fig. S2B). Interestingly, the papillary tumors appear to exhibit two distinct morphological subtypes. The Type I subtype is primarily composed of round-shaped cells of relatively uniform size, and the stratification of cell layers is not well defined (Fig. 2C). The Type II subtype is characterized by more pronounced stratification and composed of elongated spindle-shaped cells, with cell size increasing and nuclear-to-cytoplasm (N/C) ratio decreasing from the basal layer towards the superficial layer (Fig. 2D). The presence of such round-shaped and spindle-shaped cells has also been reported in human bladder papillary tumors (Montironi et al., 2008), and both papillary tumor subtypes persisted 6 months post electroporation (Fig. 2E).

**Figure 2.**
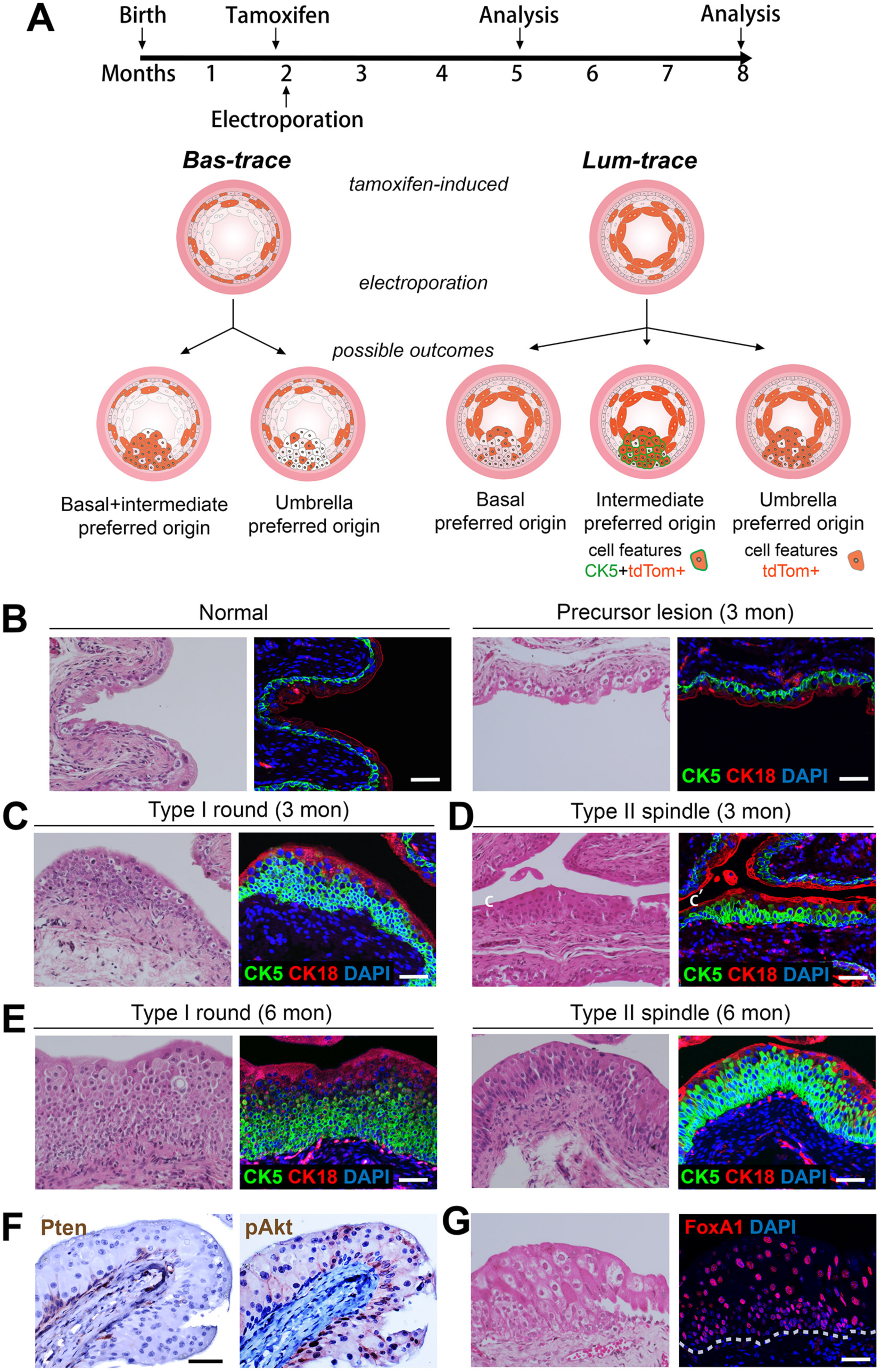
Characterization of early-stage CRISPR-induced *Pten-p53*-mutant mouse bladder tumors. (**A**) Schematic diagram showing experimental design of tracing basal and luminal cells during cancer initiation and interpretation of possible outcomes. (**B**) Adjacent sections showing H&E (left) and IF (right) images of normal and 3-month precursor lesions in the urothelium. (**C-E**) Adjacent sections showing H&E (left) and IF (right) images of representative Type I and Type II tumor subtypes at 3-month (**C**, **D**) and 6-month (**E**) post tumor induction. (**F**) Adjacent sections of IF images showing lack of Pten (left) and elevated pAkt (right) expressions in a 3-month tumor region. (**G**) Adjacent sections showing H&E (left) and IF of Foxa1 expression (right) in a 3-month tumor region. Dashed line marks the basal lamina. Scale bars in **B-G** correspond to 50 µm.

To further characterize the tumors, we next conducted immunofluorescence (IF) staining on adjacent tissue sections. Notably, both Type I and Type II tumor cells displayed enhanced CK5 expression compared to the normal urothelium, and these cells were usually surrounded by CK5-CK18+ umbrella cells on the apical side (Fig. 2C, D). The increased volume of Type I tumors appeared to result from over-proliferation of round-shaped CK5-enhanced cells, as the apical CK5-CK18+ cell layer remained thin (Fig. 2C). The increased volume for Type II tumors was contributed by both CK5-enhanced cell aggregation in the inner layers and enlarged CK5-CK18+ cell sizes in the upper layer (Fig. 2D). Loss of Pten signals and increase of pAkt signals were observed for all the tumor lesions (Fig. 2F), while FoxA1, a marker for the luminal papillary subtype (Guo et al., 2024), was strongly expressed throughout the tumor (Fig. 2G). Overall, these data demonstrate that our strategy can reliably generate bladder papillary tumors by 3 months post tumor induction. The predominance of CK5-enhanced cells in most tumors implied that inner-layer cells might have a competitive advantage over superficial-layer cells during tumorigenesis.

### Lineage analysis reveals the relative competitiveness of different urothelial cells and differentiation hierarchy

To definitively determine the capabilities of different cell types in serving as cell of origin for bladder cancer, we next examined tdTomato expression in tumors generated from both the *Bas-trace* and *Lum-trace* models. At 3 months post-electroporation, transformed urothelial cells in the inner layers were largely tdTomato-negative in the *Lum-trace* model but tdTomato-positive in the *Bas-trace* model (Fig. 3A). In contrast, transformed cells in the superficial layers adopted an inflated morphology, and were tdTomato-positive in the *Lum-trace* model but mostly tdTomato-negative in the *Bas-trace* model (Fig. 3A). These results demonstrate that both basal and luminal cells can be transformed. Next, we quantified the proportions of tdTomato-labeled or -unlabeled cells with different marker expressions out of the total urothelial cells through time. We observed that, transformed superficial-layer cells appeared to be progressively outcompeted by cells derived from inner layers. In the *Lum-trace* model, from 3 months to 6 months post-electroporation, we noticed a significant reduction of labeled umbrella tumor cells (CK5-tdTom+) (Fig. 3B, E). Those cells were displaced by expanding tdTomato-negative tumor cells (Fig. 3B, E), which originated from unlabeled basal and intermediate cells, as well as by expanding CK5+tdTom+ tumor cells (Fig. 3C, E), which originated from labeled intermediate cells. Displacement of umbrella cells was also corroborated in the *Bas-trace* model, in which CK5+tdTom+ tumor cell clusters expanded in size from 3 months to 6 months while unlabeled CK5-tdTom-umbrella cell clusters gradually shrank (Fig. 3D, F). In fact, in both models throughout progression, we never found tumor regions consisting solely of CK5-CK18+ umbrella cells. Taken together, these results suggest that although basal, intermediate, and umbrella cells could all be transformed, superficial umbrella cells were out-competed by CK5-expressing basal and intermediate cells during tumorigenesis.

**Figure 3.**
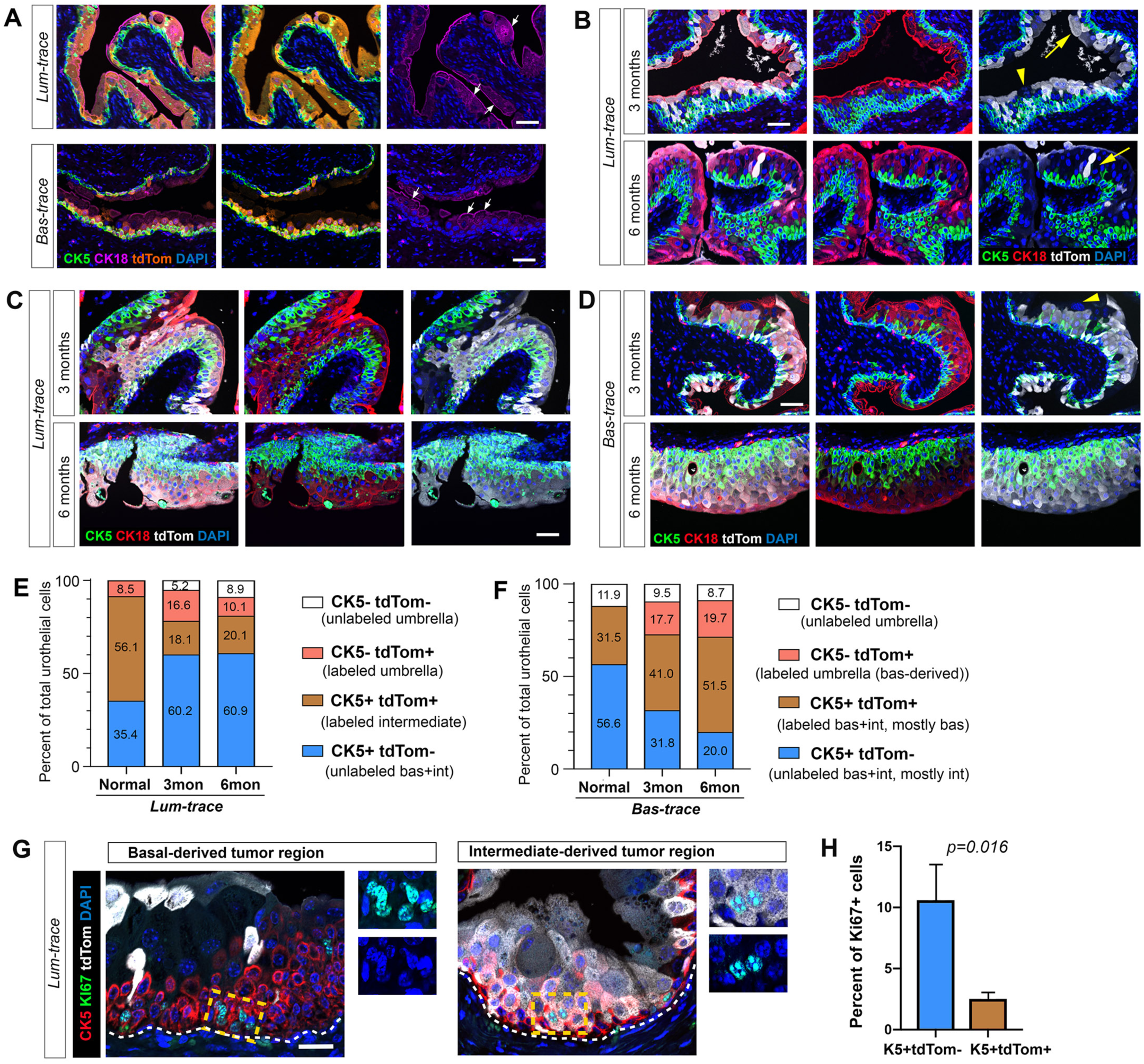
Lineage-tracing analysis comparing the competitiveness of different urothelial cell types in serving as cell of origin. (**A**) Representative IF images of 3-month tumor regions showing labeling of umbrella and intermediate cells in the *Lum-trace* mice (upper) and labeling of basal and intermediate cells in the *Bas-trace* mice (lower). Arrows pointing to enlarged umbrella cells after transformation. (**B**) Representative IF images showing decrease of labeled umbrella cells from 3-month to 6-month in the *Lum-trace* mice. Arrows pointing to labeled umbrella cells. Arrowhead pointing to unlabeled umbrella cells, which were basal/intermediate-derived. (**C**) Representative IF images showing tumor regions of labeled intermediate cells in the *Lum-trace* mice. (**D**) Representative IF images showing expansion of labeled basal cells and their luminal differentiation from 3-month to 6-month in the *Bas-trace* mice. Arrowhead pointing to a transformed unlabeled umbrella cell. (**E, F**) Bar graphs quantifying the percentages of CK5-tdTom-, CK5-tdTom+, CK5+tdTom+, and CK5+tdTom-cells out of total urothelial cells through time in the *Lum-trace* (**E**) and *Bas-trace* (**F**) mice. (**G**) IF images showing basal-derived tumor region (CK5+tdTom-) contained more Ki67+ cells than intermediate-derived tumor region (CK5+tdTom+) in the *Lum-trace* mice. (**H**) Bar graph quantifying the result of **G**. N=3 animals analyzed. Errors bars correspond to one s.d., calculated by student’s *t*-test. Scale bars in **A-D, G** correspond to 50 µm.

Quantitation of cell clusters of different markers also suggests basal cells being more aggressive than intermediate cells in bladder cancer initiation. As shown in Fig. 3E, in the *Lum-trace* model, CK5-tdTom+ cells (the entire umbrella cells) represented 8.5% of the total urothelial cells before tumor induction. By 3 months post-induction, its proportion increased to 16.6%. Since we have established that umbrella cells were not competitive in tumorigenesis, such expansion can only be explained by CK5+tdTom+ intermediate cells differentiating into CK5-tdTom+ umbrella cells during tumor initiation. Notably, by 6 months post-induction, its proportion decreased back to 10.1%, suggesting that unlabeled basal-derived tumor cells expanded even faster than intermediate-derived tumor cells. Indeed, labeled intermediate cells (CK5+tdTom+ cells) represented 56.1% of total cells before tumor induction, but shrank to 18.1% by 3 months, as a result of both umbrella cell differentiation and basal tumor cell expansion; while the proportion of unlabeled CK5+tdTom-cells (mostly basal cells plus some intermediate cells) increased significantly from 35.4% to 60.2% in the 3 months period (Fig. 3E). Furthermore, CK5-tdTom-cells, which were non-existent before tumor induction, increased to 5.2% of total cells by 3 months and further increased to 8.9% by 6 months (Fig. 3E), suggesting a continuous differentiation of unlabeled basal cells towards intermediate and umbrella cells.

Our conclusion is also supported by data obtained from the *Bas-trace* model. We observed a continuous expansion of the proportion of CK5+tdTom+ cells but shrinkage of CK5+tdTom-cells throughout tumor progression in *Bas-trace* mice (Fig. 3F). Since unlabeled CK5+tdTom-cells were initially mostly intermediate cells rather than basal cells (Fig. 1F), the relative shrinkage of this population through time is consistent with the notion that transformed intermediate cells were less proliferative than basal cells. Indeed, when we performed Ki67 IF staining in the *Lum-trace* model at 3-month post tumor induction, we observed higher proliferation in CK5+tdTom-cells, which were mostly basal-derived tumor cells, compared to CK5+tdTom+ cells, which were strictly intermediate-derived tumor cells (Fig. 3G, H). Overall, these data indicate the competitiveness of different cell types in serving as the cell of origin for bladder cancer in the following order: basal > intermediate > umbrella, which resembles the lineage hierarchy during bladder organogenesis (Kim et al., 2020). Notably, due to their deeper positioning, basal cells are less accessible than umbrella cells to CRISPR plasmids during electroporation (Fig. 1C), which makes our findings even more striking.

### Basal cells produce more heterogeneous tumor phenotypes and are the source of aggressive tumor sub-cell types

Having established the cell lineage hierarchy in bladder tumor initiation, we next sought to compare later-stage tumors originated from different cell types at both the pathological and molecular levels, and characterize tumor cell behaviors more closely. To this end, we established cell-type-specific inducible CRISPR bladder cancer models. To produce umbrella/intermediate-cell-derived tumors, we generated *Upk2-CreER^T2^; R26-LSL-Cas9-eGFP* mice (Platt et al., 2014) (referred to as the *Lum-induce* model), in which Cas9 expression is activated by the Upk2 promoter upon tamoxifen induction, ensuring that tumors arise exclusively from Upk2+ cells after electroporation of the *pCAG-eCFP-dual-sgPten-dual-sgTrp53* plasmid and that tumor cells are marked by GFP. Similarly, to produce basal/intermediate-cell-derived tumors, *CK5-CreER^T2^; R26-LSL-Cas9-eGFP* mice (referred to as the *Bas-induce* model) were used (Fig. 4A). Two weeks after tamoxifen induction, we confirmed eGFP expression in respective cell types in the two models. We then examined bladder tumor phenotypes at 8- and 13-month post-electroporation, at which time points the umbrella-derived tumor cells should have already been out-competed based on previous lineage tracing data. Therefore the *Lum-induce* tumors were mostly derived from intermediate cells, whereas the *Bas-induce* tumors were derived from both basal and intermediate cells.

**Figure 4.**
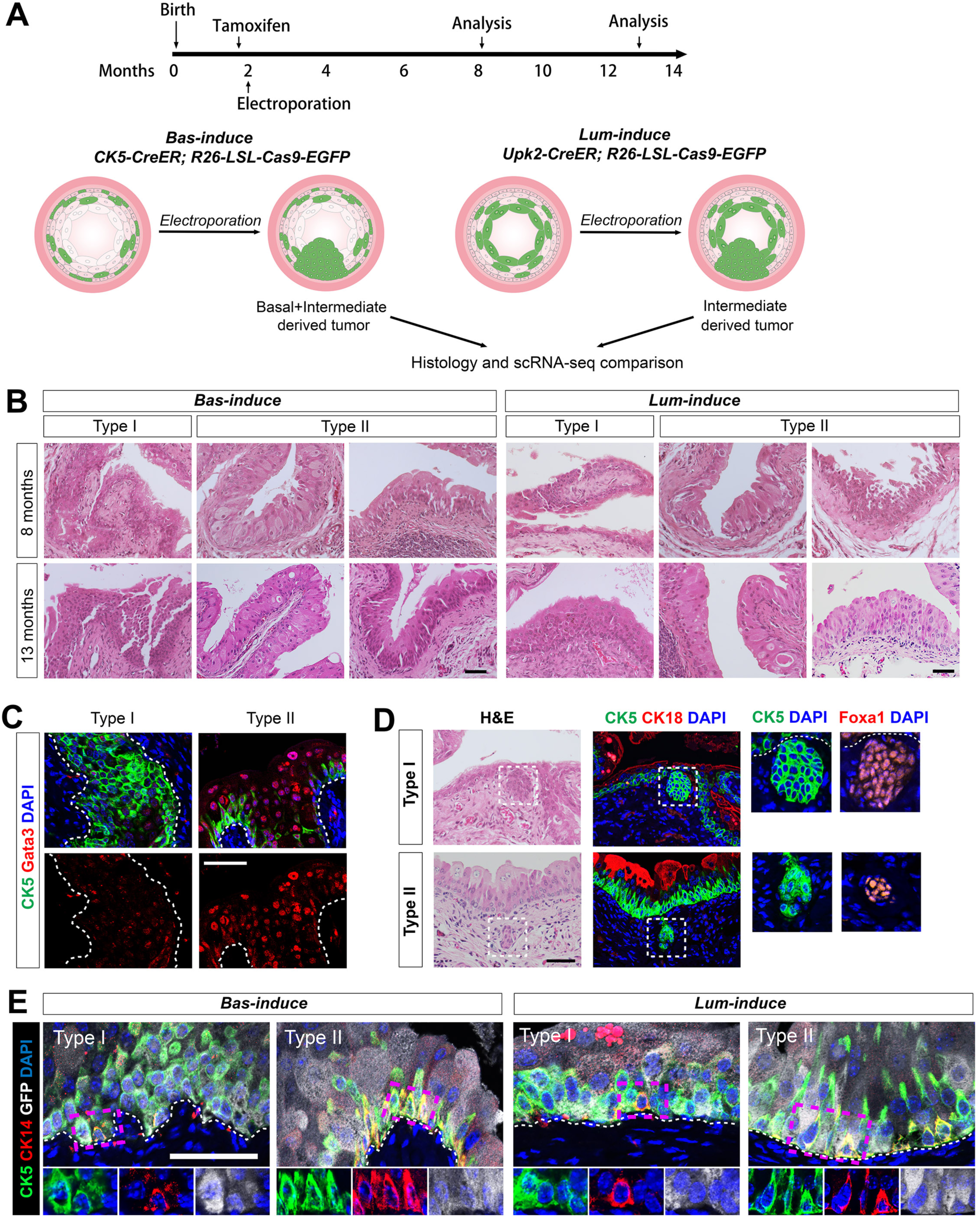
Histological characterization of *Bas-induce* and *Lum-induce* tumors. (**A**) Diagram showing experimental timeline and strategy for generating and comparing *Bas-induce* and *Lum-induce* tumors. (**B**) Representative H&E images showing Type I and Type II tumor regions in Bas-induce and Lum-induce mice at 8-month and 13-month post-electroporation. 13-month Type I tumors show CIS features. (**C**) IF staining showing higher Gata3 expression in Type II cells than Type I cells. (**D**) Adjacent sections of H&E images (left) and IF staining (middle and right zoom-in) showing the micro invasions express CK5 and Foxa1, but not CK18 in both Type I (upper row) and Type II (lower row) tumors. (**E**) IF staining showing the presence of CK14+ cells in a subset of CK5+ cells in both Type I and Type II tumors of *Bas-induce* and *Lum-induce* mice. Zoom-in view of the insets at the bottom. Dashed lines in **C-E** mark the basal lamina. Scale bars in **B-E** correspond to 50 µm.

At 8-month post-electroporation, both *Lum-induce* and *Bas-induce* models reliably produced papillary tumors that contained both round-shaped Type I and spindle-like Type II phenotypes observed in the *Lum-trace* and *Bas-trace* models (Fig. 4B). We surveyed luminal markers of human bladder cancer (Guo et al., 2024), and found that Gata3 expression was preferentially enhanced in Type II tumors compared to Type I (Fig. 4C), offering a marker for distinguishing the two subtypes. Notably, we identified occasional micro-invasions into the lamina propria underneath both Type I and II tumors (Fig. 4D). These invasive cells were Foxa1+ and CK5+ (Fig. 4D), suggesting they had intermediate cell features. We also observed that in both Type I and Type II tumors of *Lum-induce* and *Bas-induce* mice, a subset of CK5+CK18+ intermediate tumor cells began to express CK14 (Fig. 4E), which is rarely expressed in the normal adult urothelium but marks a subpopulation of basal cells associated with higher self-renewal and regeneration capabilities (Volkmer et al., 2012; Papafotiou et al., 2016), indicating that these cells might have adopted more stem-like properties.

At 13-months post-electroporation, the aforementioned papillary tumor phenotypes persisted in both models and some developed into CIS (Fig. 4B). Large GFP-positive tumor clones consisting of mostly CK5-CK18+ umbrella cells and a few CK5+ basal/intermediate cells near the basement were frequently observed in both models (Fig. 5A), confirming that both transformed basal and intermediate cells can produce umbrella tumor cells. These regions resemble the luminal CIS features in human (Wullweber et al., 2021). However, despite these similarities, compared to *Lum-induce* tumors, *Bas-induce* tumors displayed greater phenotypic diversity. A few mice in the *Bas-induce* cohort developed very large tumors, in which the majority of cells were CK5+CK18-, with a few CK5-CK18+ cells scattered within the tumor mass (Fig. 5B). Moreover, the proportion of CK14+ cells within the CK5+ cell population increased dramatically in *Bas-induce* tumors at 13-month post-electroporation (Fig. 5C), and was much higher than that in *Lum-induce* tumors (Fig. 5D). Many of these cells express Ki67 at much higher rate than CK5+CK14-cells (Fig. 5E, F), indicating that the CK5+CK14+Ki67+ triple-positive cells were the primary driver of tumor growth. In contrast, such triple-positive cells were rarely found in the *Lum-induce* tumors. The weak expressions of FoxA1 and Gata3 in these large tumors (Fig. 5G, H) were consistent with their basal tumor features, suggesting they resemble the basal pathological subtype of human bladder cancer. Indeed, these large tumors often contained regions of squamous differentiation (Guo et al., 2024) (Fig. 5I), which was not observed in the *Lum-induce* tumors.

**Figure 5.**
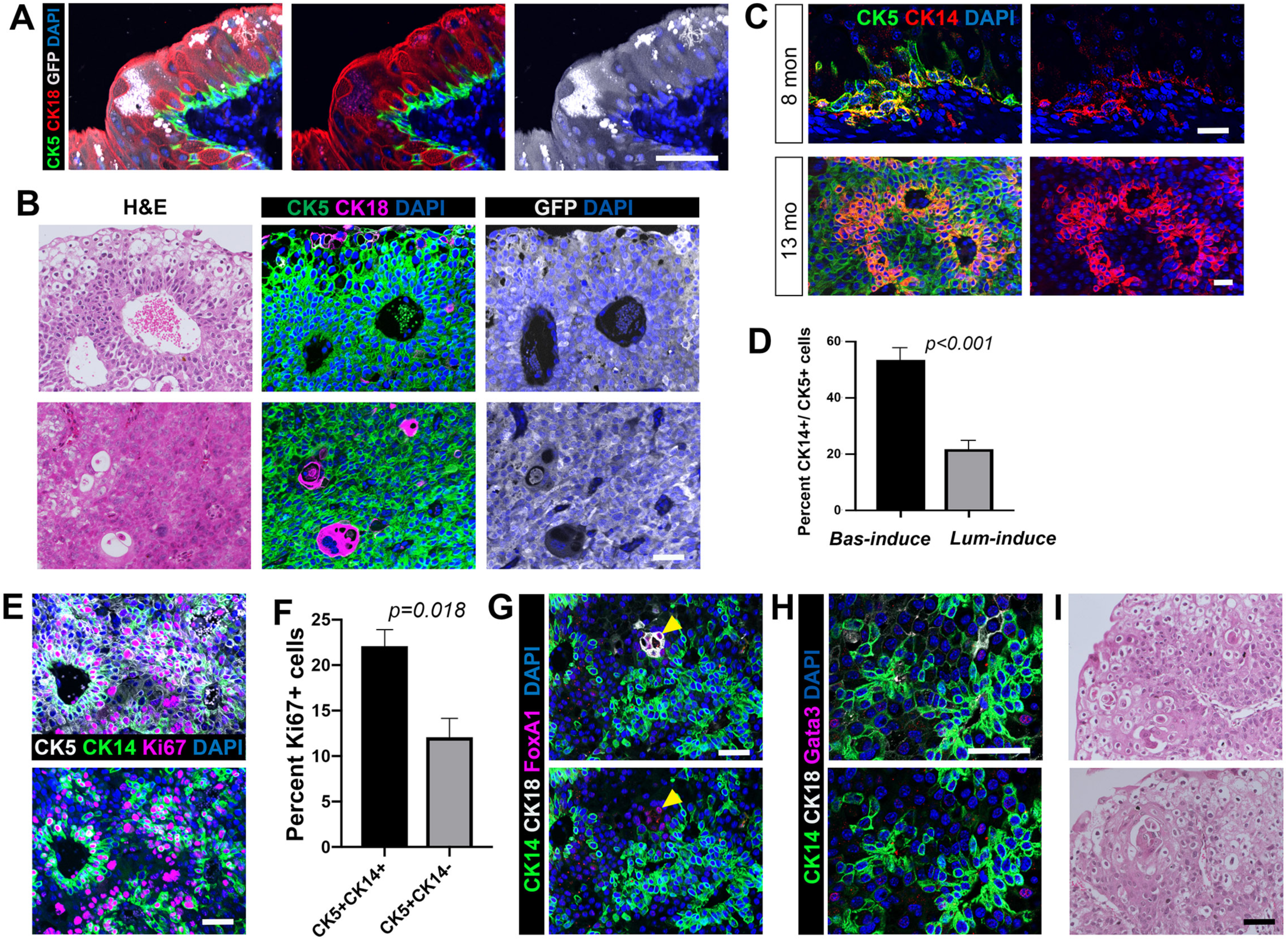
*Bas-induce* tumors have unique histological features that resemble more aggressive bladder cancer. (**A**) Representative image showing a large GFP+ tumor clone consisting mostly of CK18+CK5-cells at 13-month post-electroporation. (**B**) Adjacent sections of H&E (left) and IF staining (middle and right) in two representative loci of extra-large *Bas-induce* tumors showing their prominent basal cell feature. (**C**) IF staining showing increased CK14+ percentage in CK5+ cells from 8-month to 13-month in *Bas-induce* tumors. (**D**) Bar graph showing higher CK14+ percentage in CK5+ cells of *Bas-induce* tumors compared to *Lum-induced* tumors. N=3 tumors analyzed. P value calculated by student’s *t*-test. (**E**) Representative IF image of a CK5+CK14+Ki67+ region in the 13-month *Bas-induce* tumor. (**F**) Bar graph showing higher Ki67 index in CK5+CK14+ than CK5+CK14-cells. N=3 tumors analyzed. P value calculated by student’s *t*-test. (**G, H**) The triple-positive regions in *Bas-induce* tumors have few Foxa1 (**G**) and Gata3 (**H**) cells. Arrowhead pointing to a rare CK18+Foxa1+ luminal locus. (**I**) Two representative H&E images showing squamous differentiation in *Bas-induce* tumors. Error bars in **D**, **F** correspond to one s.d. Scale bars in **A-C, E, G-I** correspond to 50 µm.

### Molecular comparison of *Bas-induce* and *Lum-induce* tumors by scRNA-seq

We next aimed to understand the cellular and molecular compositions of *Bas-induce* and *Lum-induce* tumors by comparing their transcriptomic profiles using single cell RNA-sequencing (scRNA-seq). Tumor samples from *Bas-induce* and *Lum-induce* mice (N=3 for each) were collected at 13-month post-electroporation. Standard 10x Genomics scRNA-seq protocol was used to generate transcriptomic profiles of 8,305 and 7,580 cells for *Bas-induce* and *Lum-induce* samples, respectively. These datasets were integrated using Harmony (Korsunsky et al., 2019) to generate 17 distinct clusters, which were broadly annotated into epithelial, fibroblast, myofibroblast, endothelial, and immune cell types based on known cell markers (Fig. 6A). Notably, epithelial cells (EPCAM+) were the most diverse population with high cell count and clusters.

**Figure 6.**
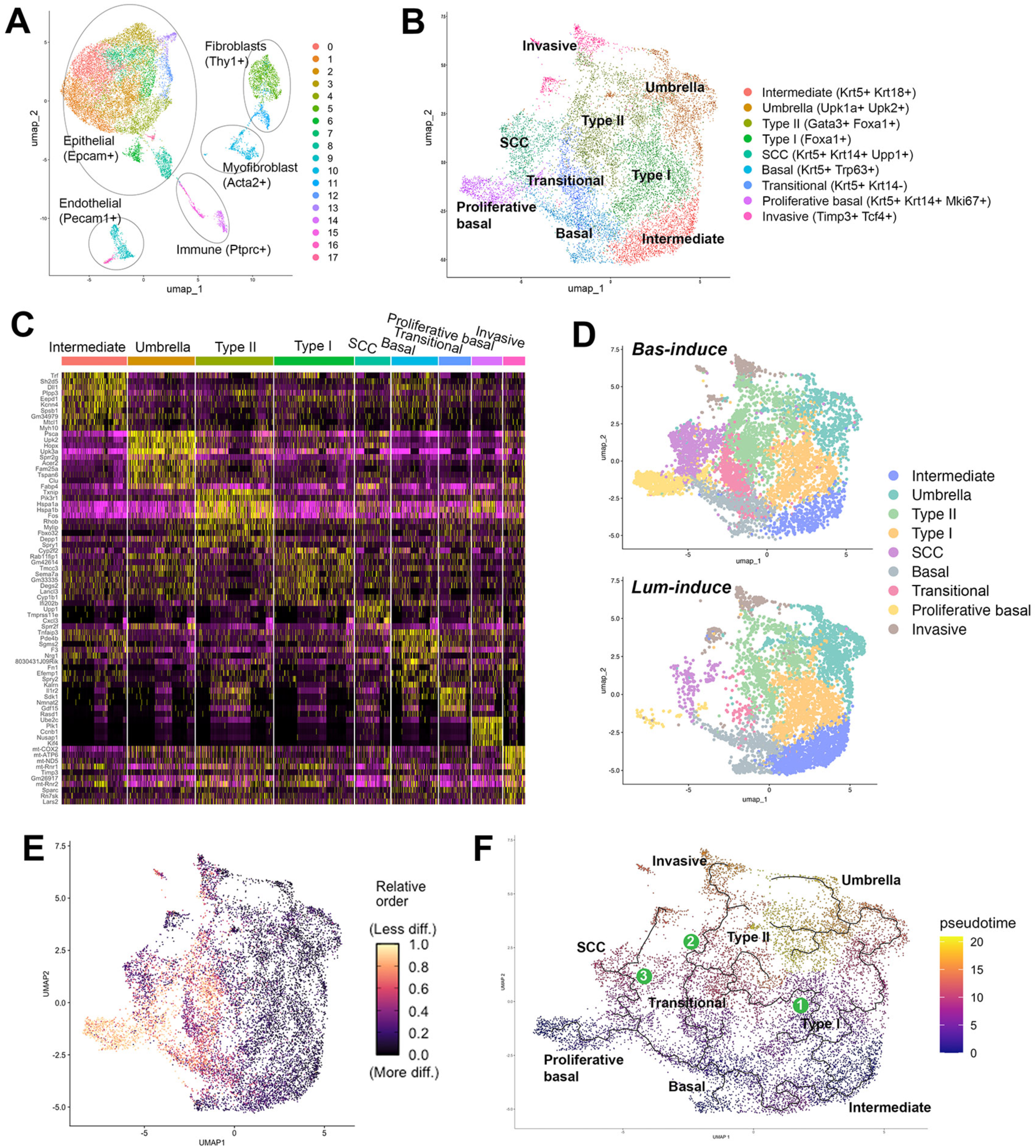
scRNA-seq analysis comparing *Bas-induce* and *Lum-induce* tumors. (**A**) UMAP showing various cell clusters and annotated cell types for all the samples. (**B**) UMAP showing different cell types (identifiable by markers shown on the right) within urothelial cells of all samples. (**C**) Heatmap showing top marker genes in each urothelial cell type. (**D**) Split UMAP view of *Bas-induce* and *Lum-induce* samples showing the overlapping cell clusters and basal-unique cell clusters. (**E**) CytoTRACE analysis showing less cell differentiation in the basal-unique cell clusters. (**F**) Pseudotime trajectory analysis showing the roots at Basal and Intermediate cells and three major differentiation lineages.

To characterize and compare the heterogeneity of the *Bas-induce* and *Lum-induce* tumors, EPCAM+ epithelial cells were re-clustered using anchor points from each cohort and harmonized to create 15 new epithelial subclusters (Fig. S3A). We then curated 9 cluster annotations encompassing both normal and tumor cell types according to expression of known marker genes and significant variable genes calculated (Fig. 6B, C). Normal cells were classified as Basal (*Krt5+, Trp63+)*, Umbrella (*Upk1a+, Upk2+*), and Intermediate (*Krt5+, Krt18+*) cell clusters. Tumor cells were distinguishable from normal based on relative high expression of oncogenic cell markers identified in previous histological comparisons. In particular, luminal tumor cells were classified as Type I (*Foxa1+*) and Type II (*Gata3+, Foxa1+*), both of which displayed modest Krt5 and Krt18 expressions (Fig. S3B). Separation of the Intermediate and Type I clusters based on *Foxa1* expression did not provide a clear divide (Fig. S3B), indicating their transcriptomic similarity and a possible gradual transformation. In agreement with histology, Type II tumors displayed higher expression of Gata3 than Type I (Fig. S3B).

Importantly, split view of UMAP based on cohorts showed that the *Bas-induce* samples contained three unique cell clusters that were rare in the *Lum-induce* samples, namely, Squamous cell carcinomas (SCC) (*Krt5*+, *Krt14*+, *Upp1*+), Proliferative basal (*Krt5*+, *Krt14*+, *Mki67*+) and Transitional (*Krt5*+, *Krt14*-) (Fig. 6D). The SCC and Proliferative basal cells have been identified in previous histology analysis (Fig. 5E, I), while the Transitional cells retain strong basal identity but also express modest levels of Gata3 (Fig. S3B), indicating transition towards Type II cells. Interestingly, CytoTRACE analysis (Gulati et al., 2020) revealed that these basal-unique clusters, particularly the Proliferative basal cells, were the most stem-like and least differentiated (Fig. 6E), implicating their aggressive nature. Lastly, a cluster we named Invasive cells (*Timp3+*, *Tcf4+*) has modest basal and luminal marker gene expressions (Fig. S3B). Marker genes for each annotated urothelial cluster is listed in Supplemental File 1.

Quantification of the proportions of different cell clusters in the two cohort samples revealed the strong bias of SCC, Proliferative basal, and Transitional, and moderate enrichment of Type II cells in *Bas-induce* samples, and a slight preference of Type I cell enrichment in *Lum-induce* samples (Fig. S3C). To determine the cellular differentiation hierarchy during tumor progression, we next employed monocle3 (Trapnell et al., 2014) to assemble a pseudotime trajectory of the urothelial cell clusters. Utilizing unsupervised selection of roots, Basal and Intermediate cell clusters were determined as points of entry, with the *Bas-induce* samples also having a unique root at the Proliferative basal cluster (Fig. 6F). From this structure we observed three distinct tumor lineages. Intermediate cells preferentially generate Type I cells, which can further progress into Type II cells. Notably, Type II cells can be traced back to both Basal and Intermediate lineages, consistent with the histology finding that spindle-shaped tumor cells were present in both *Bas-induce* and *Lum-induce* tumors. While both lineages share an upregulation of Gata3 that defines Type II tumors, lineages from the different roots display their respective marker genes in differing parts of the Type II cluster. Basal-derived Type II cells display increased *Krt14* expression whereas Intermediate-derived ones display elevated *Krt18* expression (Fig. S3D). The trajectory of Transitional cells goes from Basal to Type II and eventually into Invasive cells, indicating that basal derived tumors are prone to generate invasive phenotypes. A third unique trajectory found in the *Bas-induce* samples saw the generation of SCC and Proliferative basal cells (Fig. 6F). These basal-specific cell clusters are separate from the rest of the shared lineages, and affirm our conclusion from histology data that basal cells, when oncogenically targeted, are capable of producing all tumor cell types, including those unique aggressive populations not found in the *Lum-induce* tumors.

### Basal-cell derived gene signature correlates with human basal molecular subtype and predicts worse patient outcome

To determine whether cells of origin may underlie the different molecular subtypes of bladder cancer, we next sought to understand gene expression heterogeneity of *Bas-induce* and *Lum-induce* tumors, and compare gene signatures obtained from mouse data to clinical patient data. We first investigated the molecular differences between basal tumor clusters (Proliferative basal, SCC, and Transitioning) and luminal tumor clusters (Type I and Type II) of the urothelial UMAP. Differential gene expression (DGE) analysis showed that 1872 genes were upregulated in the basal tumor clusters and 1298 upregulated in the luminal tumor clusters (Adjusted P-value < 0.05 and log2FoldChange (FC) > 0.5) (Fig. 7A). Upregulated genes in basal tumor clusters included *Ube2c* and *Nusap1*, which are associated with aggressive bladder cancers (Morikawa et al., 2013; Chen et al., 2021; Hou et al., 2022), while upregulated genes in luminal tumor clusters are related to cell growth and metabolism such as *Enho* and *Cyp1a1* (Fig. 7A). To further understand the heterogeneity within the basal cell clusters, we performed DGE analysis comparing the Proliferative basal cluster and the SCC cluster, and found that 1766 genes were upregulated in the Proliferative basal cluster while 614 genes upregulated in the SCC cluster (Adjusted P-value < 0.05 and log2FoldChange (FC) > 0.5) (Fig. 7B). Many genes associated with aggressiveness such as Ube2c, Nusap1, and Top2a appeared in the upregulated gene list of the Proliferative basal cluster (Fig. 7B), indicating these highly proliferative basal cells being the most aggressive population in the *Bas-induce* tumors. Next, we performed gene set enrichment analysis (GSEA) (Subramanian et al., 2005) to compare luminal and basal tumor clusters using the genes found in the DGE analysis. Consistently, basal tumor clusters showed a high enrichment in cell cycle G2M checkpoint and Myc targets (Fig. 7C), underlying their more proliferative nature, while luminal tumor clusters were enriched for Notch signaling and xenobiotic metabolism (Fig. 7D), the latter of which corroborates with a report showing that some luminal subtypes of bladder cancer patients exhibit resistance to cisplatin-based chemotherapy (Choi et al., 2014b).

**Figure 7.**
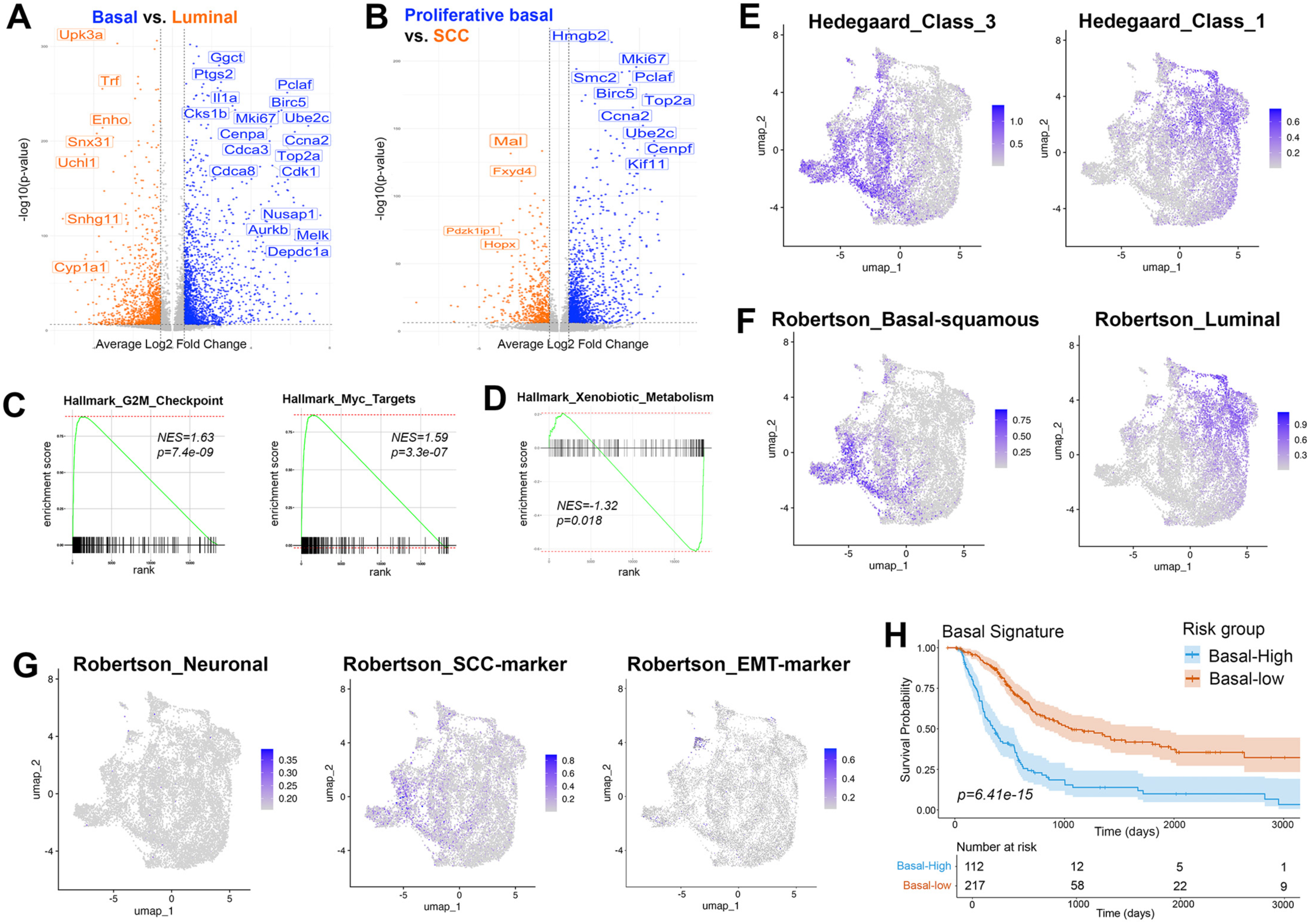
Basal-cell derived gene signature correlates with human basal molecular subtype and predicts worse patient outcome. **(A)** Volcano plot showing DGE of basal vs. luminal tumor cell clusters. (**B**) Volcano plot showing DGE of Proliferative basal vs. SCC tumor cell clusters. (**C, D**) GSEA showing enrichment of pathways in basal tumor clusters (**C**, positive NES) vs. luminal tumor clusters (**D**, negative NES). (**E-G**) Feature plots depicting molecular subtype classifiers reported in the Hedegaard (**E**) and Robertson (**F, G**) datasets are enriched in corresponding mouse tumor cell clusters. (**H**) Kaplan-Meier curve on TCGA-BLCA cohort using the unique basal signature showing patients with high basal signature having worse overall survival.

Next, we performed mouse-human cross-species comparison by interrogating two published human datasets of bladder molecular subtypes against our scRNA-seq data. The Hedegaard dataset (Hedegaard et al., 2016) profiled the transcriptional landscape of NMIBC and identified three molecular subtypes that displayed luminal and basal characteristics. We utilized these classifiers to associate these molecular properties with our mouse tumor clusters. We found that Class_3 signature, which was defined as human basal-like subtype, showed strong enrichment in our mouse basal unique clusters (Fig. 7E). Conversely, Class_1, which was the less aggressive human luminal-like subtype, showed strong signature enrichment in the mouse luminal tumor cell clusters such as Type I and Type II (Fig. 7E). Moreover, the Robertson dataset (Robertson et al., 2017) defined 5 molecular subtypes from MIBC: basal_squamous, luminal, luminal papillary, luminal-infiltrated, and neuronal. We found that the enrichment score for the basal_squamous and luminal subtypes again correlated well with the corresponding mouse tumor clusters in our UMAP (Fig. 7F). No neuronal score enrichment was found (Fig. 7G), likely due to the inability of our mouse models in generating neuroendocrine tumors. However, a SCC_marker human signature and an EMT_marker human signature used in the Robertson paper overlapped well with our mouse SCC cluster and the Invasive cluster in the UMAP, respectively (Fig. 7G). Together, these data strongly support that the different molecular subtypes of human bladder cancer have the corresponding cells of origin as their basis.

Finally, we generated a molecular gene signature for our unique basal tumors and compared it to patient data taken from the TCGA-BLCA cohort (n = 330) (Cancer Genome Atlas Research, 2014). This basal molecular gene signature was constructed from the list of upregulated genes identified in DGE in our Proliferative Basal, SCC and Transitional clusters against all other clusters. These genes were screened using a univariate Cox regression to remove non-informative genes (p > 0.05) from the signature. 87 genes were associated with our basal tumors, which were used to construct a multivariate Cox proportional hazard model (C = 0.76; 95% CI 0.73–0.80; P = 4.93e-06) (Supplemental File 2) and were used to stratify patients into the Basal-High and Basal-Low groups. Kaplan-Meier curves were then constructed using the risk associated with basal tumors (log-rank P = 6.41e-15) (Fig. 7H). Curves show significant separation between the two cohorts, indicating the strong correlation of our basal unique signatures with poor overall survival.

## Discussion

Despite significant progress in the molecular classification of bladder cancer, the identity and contribution of distinct urothelial cell types to tumor initiation remain incompletely understood. Previous studies have suggested that basal and intermediate cells may give rise to histologically and molecularly distinct tumors (Shin et al., 2014; Van Batavia et al., 2014; Papafotiou et al., 2016; Tate et al., 2021), but these insights have largely relied on chemical carcinogenesis models, such as BBN, which introduce uncontrolled mutational variability and confound interpretations about intrinsic cellular susceptibility. Moreover, it remains unclear whether the molecular subtypes of bladder cancer truly reflect their cells of origin, or if they emerge secondarily through tumor evolution and microenvironmental influences. Resolving these uncertainties is critical for understanding bladder cancer pathogenesis and for the development of cell-type-specific therapeutic strategies.

Our study establishes a novel CRISPR/Cas9-based, cell-type-specific lineage tracing system to interrogate the cell of origin in bladder cancer under defined genetic contexts. By introducing defined oncogenic events (*Pten* and *Trp53* inactivation) into specific urothelial compartments, our approach enables a direct comparison of the tumorigenic potential of basal, intermediate, and umbrella cells. Although intermediate-cell-specific CreER driver is currently unavailable, we were able to determine intermediate cell properties by carefully analyzing the combined lineage-tracing data obtained from *CK5-CreER* and *Upk2-CreER* drivers, since each driver cleanly excludes the cell layer at the opposite end (no umbrella marking by *CK5-CreER* and no basal marking by *Upk2-CreER*). Our results show that all three major urothelial cell types can serve as cells of origin for bladder cancer, yet basal cells exhibit markedly higher competitiveness, generating tumors with greater proliferative capacity, phenotypic diversity, and molecular heterogeneity. Notably, tumors arising from basal cells harbor unique proliferative basal (Krt5+Krt14+Ki67+), transitional, and squamous-like populations, which were rare in tumors arising from intermediate cells. Trajectory analysis and histological progression further revealed that basal-derived tumors can give rise to both basal-like and luminal-like lineages, supporting a model in which basal cells sit atop a tumor differentiation hierarchy. These results are consistent with the behavior of basal cells during injury repair and organoid formation (Lee et al., 2018; Kim et al., 2020; Wiessner et al., 2022), where they exhibit the broadest differentiation potential.

Our findings also for the first time directly link the molecular subtypes of bladder cancer to their corresponding cells of origin. scRNA-seq data revealed that mouse tumors derived from basal cells are enriched in gene expression signatures corresponding to human basal-squamous subtypes, whereas intermediate-derived tumors more closely resemble luminal subtypes. Furthermore, a gene signature derived from aggressive basal-specific tumor clusters in mice predicts poorer prognosis in human patients, supporting the clinical relevance of cell-of-origin–based classification. These observations provide strong evidence that the cell of origin contributes not only to the initiation of bladder cancer but also to the trajectory of its molecular evolution.

Interestingly, the prominent potency of basal cells in bladder cancer initiation contrast with our prior findings obtained through similar approaches in prostate cancer, where luminal cells, rather than basal cells, are more susceptible to transformation and serve as the predominant cell of origin under similar *Pten* inactivation context (Wang et al., 2013; Wang et al., 2014). This highlights tissue-specific differences in epithelial hierarchies and plasticity. Whereas prostate basal cells are more resistant to oncogenic reprogramming *in vivo*, bladder basal cells appear inherently more plastic and tumorigenic, possibly due to differences in stem cell niches, chromatin accessibility, or developmental programming. Our findings thus emphasize that the relationship between cell of origin and tumor subtype is organ-specific and cannot be generalized across epithelial cancers.

Notably, we observed a bias in our CRISPR targeting efficiency, with superficial umbrella cells being more readily transfected than deeper basal cells. This makes the dominance of basal-derived tumors all the more striking but also raises the possibility that rare umbrella-derived tumors might be underrepresented. Additionally, although our model recapitulates multiple key features of human bladder cancer, including subtype diversity and squamous differentiation, it does not fully reproduce the entire spectrum of human bladder cancer phenotypes due to the limitations of the *Pten/Trp53*-specific mutant background. The relative influence of different cells of origin versus different genetic mutant backgrounds on bladder cancer progression remains to be tested in the future. Nevertheless, our work for the first time provides direct experimental evidence that the molecular subtypes of bladder cancer can be traced back to the corresponding urothelial cell types, with basal cells possessing intrinsic properties that favor the development of more aggressive, heterogeneous tumors. By elucidating the cellular roots of bladder cancer subtypes, our study lays a foundation for rational design of therapies targeting cell-of-origin–specific vulnerabilities.

## Experimental Procedures

### Corresponding author

Further information and requests for resources and reagents should be directed to and will be fulfilled by the corresponding author, Zhu A. Wang.

### Data and code availability

All data reported in this manuscript will be shared by the corresponding author upon request. The bladder scRNA-seq data will be deposited in the Gene Expression Omnibus database. This paper does not report original code. Any additional information required to reanalyze the data reported in this manuscript is available from the corresponding author upon request.

## Method Details

### Mouse strains and genotyping

The *B6.CBA-Tg(Upk2-icre/ERT2)1Ccc/J* (named *Upk2-CreER^T2^*) (Shen et al., 2012), *FVB.Cg-Tg(Krt5-cre/ERT2)2LPC/JeldJ* (named *CK5-CreER^T2^*) (Indra et al., 1999), *B6.Cg-Gt(ROSA)26Sortm9(CAG-tdTomato)Hze/J* (named *Ai9*) (Madisen et al., 2010), *B6J.129(Cg)-Gt(ROSA)26Sortm1.1(CAG-cas9*,-EGFP)Fezh/J* (named *R26-Cas9-EGFP*) (Platt et al., 2014), and *B6J.129(B6N)-Gt(ROSA)26Sor<tm1(CAG-cas9*,- EGFP)Fezh>/J* (named *R26-LSL-Cas9-EGFP*) (Platt et al., 2014) mice were obtained from JAX and maintained in C57BL/6N or mixed background. Genotyping was performed by PCR using tail genomic DNA, with the primer sequences listed in Table S1.

### Mouse procedures

For tamoxifen induction, mice were administered 9 mg per 40 g body weight tamoxifen (Sigma) suspended in corn oil by oral gavage once daily for 4 consecutive days.

For plasmid electroporation, all procedures were performed as described previously (Yu et al., 2018) with modifications. Briefly, mice were anesthetized by isoflurane and restrained on a warm plate. 20 µl of 1 mg/µl plasmid mixed with trypan blue was injected through urethra using Exel Safelet Catheter (model 24G X3/4) into the PBS pre-rinsed bladder. The plasmid was retained in the bladder by tying up the external urethral orifice after catheter removal. An incision was then made in the abdomen and the bladder was pulled out from the body by tweezers. Electroporation was carried out using two electrodes placed at opposite sides of the bladder. The electrodes were connected to an electroporation generator using the parameters of 33V, 50 ms working time, and 950 ms interval time. The bladder was placed back in the body cavity and the incision was sutured. The wellbeing of the mice was monitored immediately after the surgery and consecutively for 7 days.

For bladder sample collection, tissues were dissected and fixed in 4% paraformaldehyde for subsequent cryo-embedding in OCT compound (Sakura), or fixed in 10% formalin followed by paraffin embedding. For bladder cell dissociation, collected tissues were minced into small clumps, then enzymatically dissociated using 0.2% Collagenase/Hyaluronidase (StemCell Technologies) in DMEM/F12 medium containing 2% FBS for 1 hour at 37 °C. This was followed by digestion with 0.25% Trypsin-EDTA (StemCell Technologies) for 10 min at 37 °C, and further dissociation by pipetting in freshly made Dispase/DNase (StemCell Technologies) solution. The resulting suspension was filtered through a 40 µm cell strainer to obtain single-cell suspensions, which were finally suspended in Hanks’ Balanced Salt Solution Modified/2% FBS.

All animal experiments received approval from the Institutional Animal Care and Use Committee at UCSC.

### Design of sgRNAs and plasmid construction

To enhance gene targeting efficiency, dual sgRNAs were designed and used for each gene. Dual sgRNAs targeting *Pten* were adopted from a previous publication (Xue et al., 2014). Dual sgRNAs for *Trp53* (Fig. S1A) were newly designed using the CRISPR/Cas9 track available on the UCSC Genome Browser (https://genome.ucsc.edu). The four sgRNAs were driven by human U6, human H1, human 7SK and mouse U6 promoters, respectively, and their arrangement is depicted in Fig. S1C. The tandemly arrayed sgRNAs were then inserted into the AseI restriction enzyme site of the *pCMV-eCFP* plasmid to create the targeting plasmid *pCMV-eCFP-dual-sgPten-dual-Trp53* (Fig. S1C). The *pCMV-eCFP* plasmid was modified from pCMV-mCherry2-C1 (Addgene Plasmid #54563) by replacing the mCherry sequence with an enhanced CFP sequence.

### Histology and immunostaining

Histology was performed on 4 µm serial sections of the bladder tissue. H&E staining was performed using standard protocols as previously described, and visualized using a Zeiss AxioImager. For IF staining, paraffin sections were deparaffinized and rehydrated using xylene, graded alcohol series, and three times of 1×PBS washing. Antigen retrieval was performed by boiling tissue for 30 min in Antigen Unmasking Solution (Vector Laboratories, Inc.), and then tissues were washed with PBST ×3 times for 5 min each and incubated in 5% horse serum blocking solution for 1 hour at room temperature. Primary antibodies in 5% horse serum were incubated overnight at 4°C. After PBST ×3 times washing, secondary antibodies (diluted 1:500 in PBST) labeled with Alexa Fluor 488, 555, or 647 (Invitrogen/Molecular Probes) were incubated for 1 hour at room temperature. Slides were mounted using Antifade Mounting Medium with DAPI (Vector Laboratories, Inc.) after PBST ×3 times washing, and images were taken on a Leica TCS SP5 spectral confocal microscope in the UCSC Microscopy Shared Facility. All primary antibodies and dilutions used are listed in Table S3.

### Lineage analysis, data quantitation and statistics

For lineage-tracing analysis, cell numbers were counted manually using confocal 40x photomicrographs across tissue sections. Urothelial basal cells were determined based on positive CK5 staining and location at the basement membrane. Umbrella cells were determined based on positive CK18 and negative CK5 staining and location at the most superficial side of the urothelium facing the lumen. Intermediate cells were determined based on positive staining for both CK5 and CK18 and location between the basal and superficial layers. Statistical analyses for IF staining images were performed using the two-sided student’s t-test. At least three biological replicates for each experiment or genotype were analyzed. The variances were similar between the groups that were being statistically compared.

### Single cell RNA sequencing

For collection of 13-month *Bas-induce* and *Lum-induce* bladder tumor cells, the urothelium was separated from the muscle layer using tweezers, and tumor lesions were identified based on GFP signal under a fluorescent microscope and excised from the epithelium with a scalpel before being transferred to a Collagenase/Hydrogenase digestion solution (F12/MEMD + 4.5%FBS + 10%C/H) to obtain single cell suspension. Approximately 16,000 dissociated cells were collected from each sample. Library preparation was performed using 10x Genomics Chromium Single Cell 3’ Solution with v3.1 chemistry following the manufacturer’s protocol (10x Genomics). The library purity, size, and quantity were validated by capillary electrophoresis using 2,100 Bioanalyzer (Agilent Technologies). The libraries were sequenced at the UC Davis Genome Center with a NovaSeq 6000 S4 instrument (Illumina) to a depth of ∼225k reads per cell.

### Single-cell data preprocessing and reduction

The quality of each library was assessed and visualized using FASTQC. Reads were then processed and mapped to the mm39 genome, including transgene transcripts for YFP and tdTomato. STARSolo was used to align with additional parameters for better comparison of raw counts with CellRanger >=v3.0.0: --clipAdapterType CellRanger4, -- outFilterScoreMin 30--soloCBmatchWLtype 1MM_multi_Nbase_pseudocounts, -- soloUMIfiltering MultiGeneUMI_CR --soloUMIdedup 1MM_CR. The generated gene expression matrices were assessed for quality control metrics in the R (v4.4.3) statistical environment using Seurat (v5.2.1) (Butler et al., 2018). Low quality cells with low features (<200) and mitochondrial counts (>10%) were excluded from analysis along with genes that appeared in less than 3 cells. Normalization and extrapolation of highly variable features to be scaled were performed using sctransform (v0.4.1) under default parameters. Principal component analysis was performed using the RunPCA function and assessed for prominent components for visualization using uniform manifold approximation projection (UMAP) reductions. Datasets were then merged using IntegrateLayers with Harmony (Korsunsky et al., 2019) to preserve transcriptional diversity while correcting for batch effect.

### Cell type annotation

Differentially expressed genes were assessed at each cell cluster to determine relative cell type identity. FindAllMarkers was used with default parameters to understand relationships between large populations of clusters. Nuanced FindMarkers derived smaller cell types within larger groups to devolve the inherent bladder heterogeneity. Cell types were annotated based on prior datasets, documented marker genes, and compared to histological and molecular analysis findings.

### Trajectory analysis

To derive a trajectory of cell states in our epithelial TME, we used the monocle3 (v1.3.7) (Trapnell et al., 2014) R package. Clusters found within our UMAP were re-clustered using the Leiden algorithm to reconstruct a k-nearest neighbor graph. Inferred pseudotime was then predicted using unsupervised root selection. The branching trajectory was overlaid on the UMAP projection, where roots were re-evaluated based on biological significance.

### Differential gene expression analysis

Pairwise comparison between two clusters or groups of clusters was conducted using the AggregateExpression function in Seurat to pseudobulk groups into comparable cohorts. Pseudobulked clusters would then be analyzed using FindMarkers to extrapolate variable genes for further assessment.

### Gene set enrichment analysis

Differential gene expression matrices were assessed for statistical significance using the Wilcoxon rank sum test to calculate p-value and auROC (Area Under the Receiver Operating Characteristic) scores. Scored features were then ranked on gene expression and significance (log2FC > 0.6, p-value < 0.05) to be then assessed for pathway analysis by fgsea (Korotkevich, et al, 2019) R package using the “HALLMARK” gene sets in Molecular Signature Database (MsigDB).

### Data collection

Patient clinical and RNA-Seq expression data were retrieved from The Cancer Genome Atlas (TCGA) via the Bioconductor package TCGAbiolinks (Colaprico et al., 2016). Patients lacking gene expression data or overall survival were not considered in this study. Gene expression for each patient was analyzed using the limma-voom pipeline to identify differentially expressed genes between patient normal and primary tumor samples. Empirical Bayes moderation was used to compute moderated t-statistics and adjust for multiple testing.

### Survival curve analysis

To correlate gene signatures from our dataset to clinical data, genes found to be upregulated in each cluster were identified through DGE and pathway analysis. A multivariate Cox proportional hazard model was constructed with these genes and assessed for only those found significant (FDR-corrected Wald-test p-value > 0.05). Patients were assessed for risk using a linear sum of each gene multiplied by the hazard coefficient for the covariate. All patient model fitting was done on maximally ranked statistics using the maxstats package.

## Supporting information

Supplemental File 1 - epithelial markers

Supplemental File 2 - basal signature

## Acknowledgements

We thank the microscopy core facilities at UCSC and the genomics core facilities at UC Davis for technical support. This work was supported by a CIRM postdoctoral training fellowship to C.Y. and a CIRM pre-doctoral training fellowship to N.C., and by NIH grant R01CA271452 and ACS grant 134386-RSG-20-038-01-DDC to Z.A.W.

## Author contributions

C.Y., N.C., and Z.A.W. designed the study. C.Y. performed mouse experiments with aid from A.A., J.G., and Q.X. N.C. performed all bioinformatic analyses. B.K. evaluated tumor pathology. All the authors discussed data. Z.A.W., C.Y., and N.C. prepared figures and wrote the manuscript.

## Supplemental Figures and Tables

**Figure S1.**
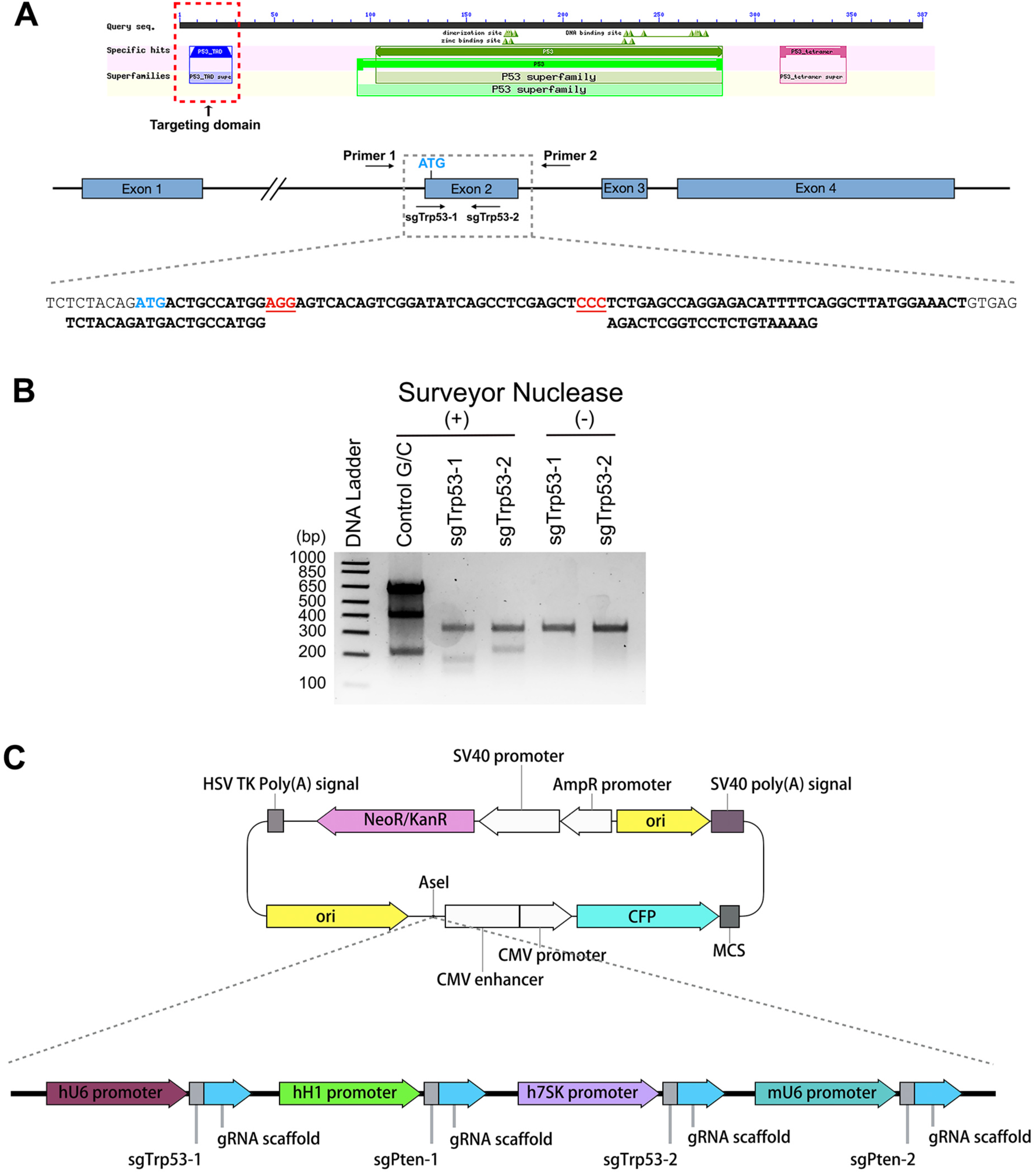
Design and validation of *Pten* and *Trp53* CRISPR targeting plasmid. (**A**) Location of the two sgRNAs targeting the *Trp53* gene locus. The two sgRNAs that we used for targeting *Pten* have been reported and validated in a previous publication (Xue et al., 2014). (**B**) Surveyor assay confirming the targeting efficacy of the two *Trp53* sgRNAs. (**C**) Diagram showing the molecular cloning strategy and structure of the *pCAG-eCFP-dual-sgPten-dual-sgTrp53* plasmid.

**Figure S2.**
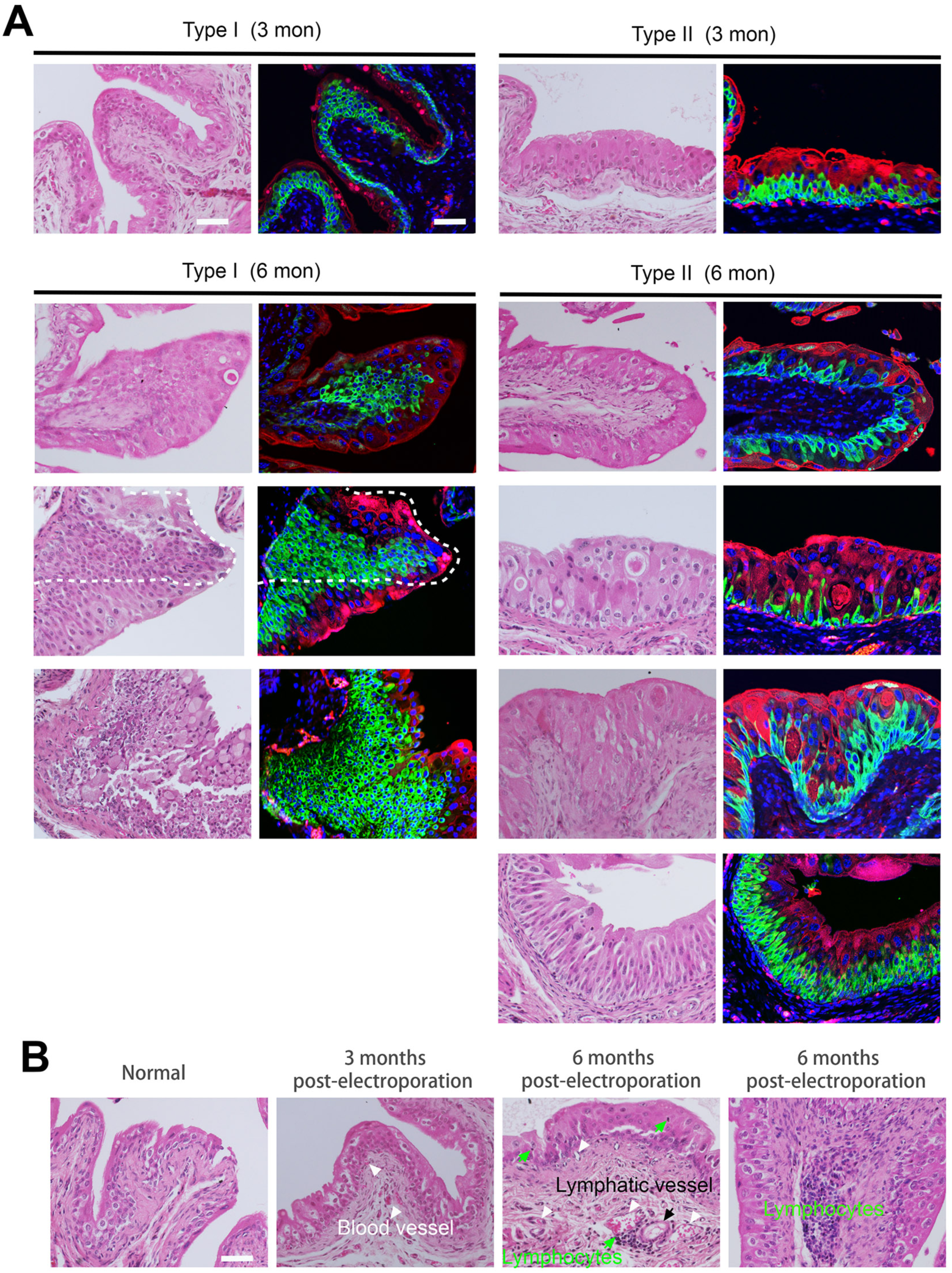
Characterization of early-stage CRISPR-induced *Pten-p53*-mutant mouse bladder tumors. (**A**) Adjacent sections showing H&E (left) and IF (right) images of representative Type I and Type II tumors at 3 months and 6 months, highlighting variations in cell size, morphology and eosinophilic staining within the same tumor type. (**B**) Representative images showing angiogenesis and lymphatic infiltration, particularly in the disorganized tumor-adjacent lamina propria in 3- and 6-month tumors. White arrowheads indicate blood vessel, black arrow indicates lymphatic vessel, and green arrows indicate lymphocytes. Scale bars correspond to 50 µm.

**Figure S3.**
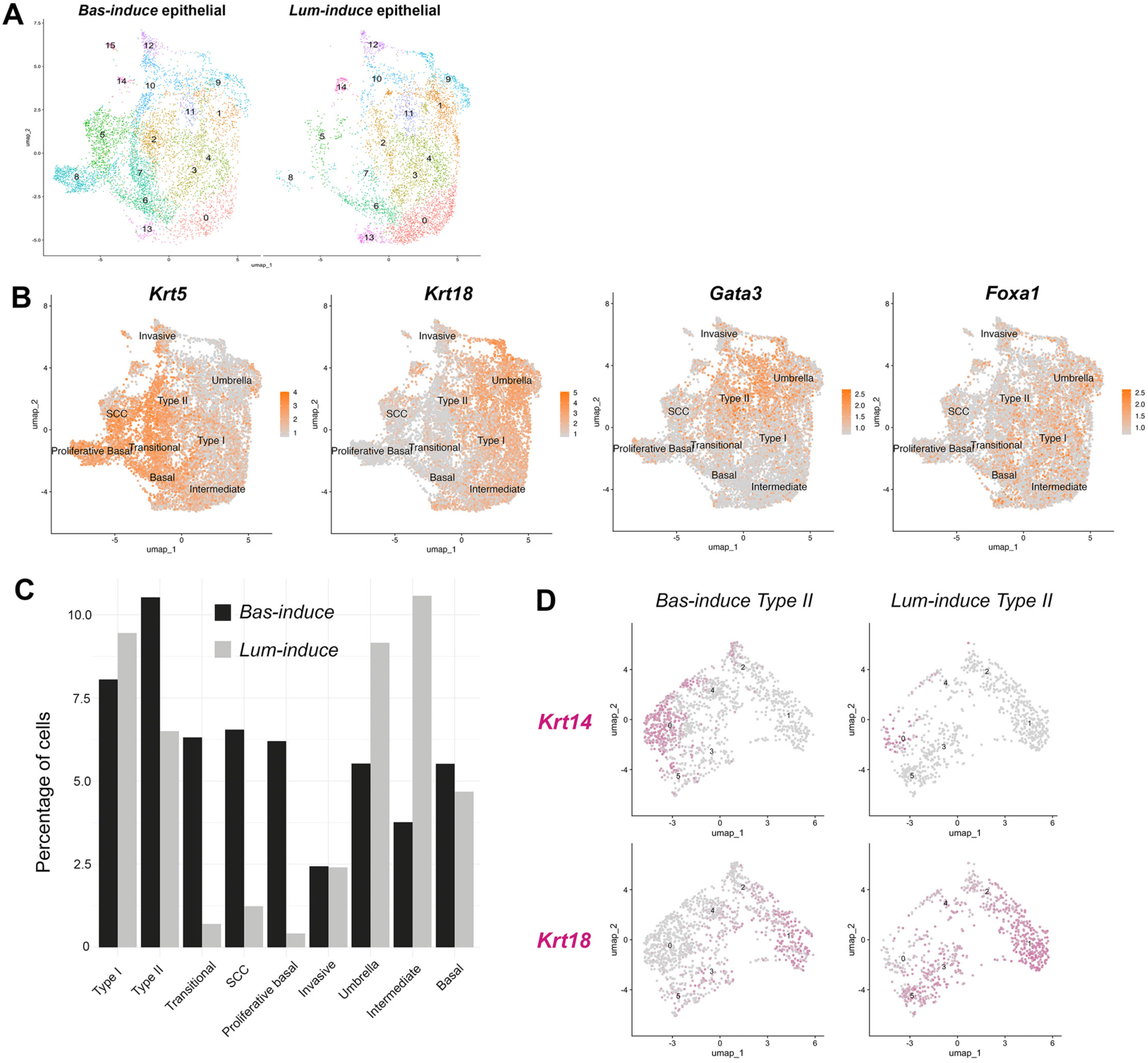
Gene expression in urothelial cell clusters of the *Bas-induce* and *Lum-induce* tumors. (**A**) Split view of UMAP of re-clustered epithelial (EPCAM+) cells in the *Bas-induce* and *Lum-induce* samples. (**B**) Feature plots of epithelial markers (Krt5, Krt18) and genes associated with bladder cancer (Gata3, Foxa1) in the UMAP of total epithelial cells. (**C**) Bar graph showing cell count percentages of the epithelial sub-clusters for the *Bas-induce* and *Lum-induce* samples. (**D**) Feature plots of *Krt14* and *Krt18* expressions in the Type II cluster of the *Bas-induce* and *Lum-induce* samples.

**Table S1.**
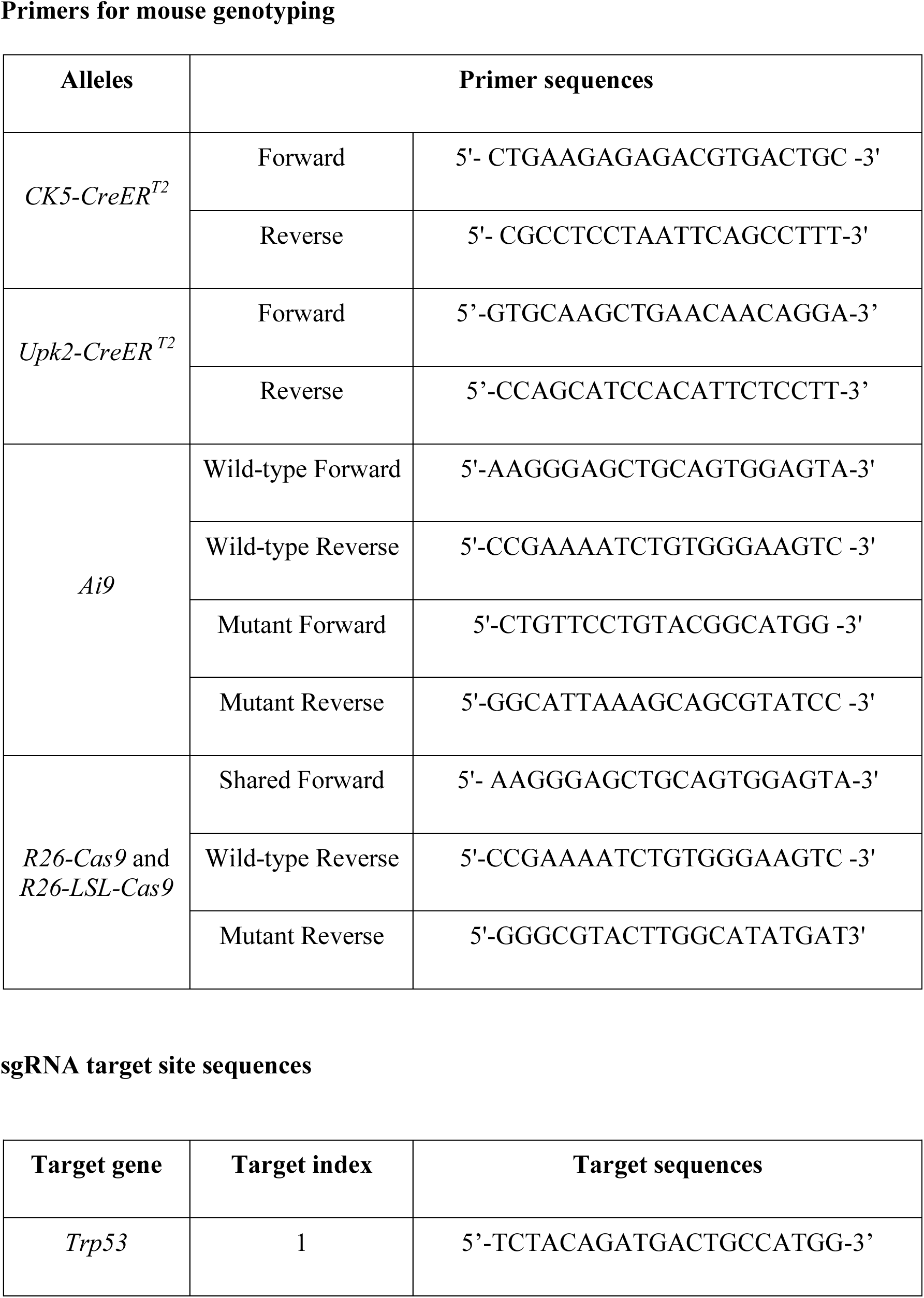
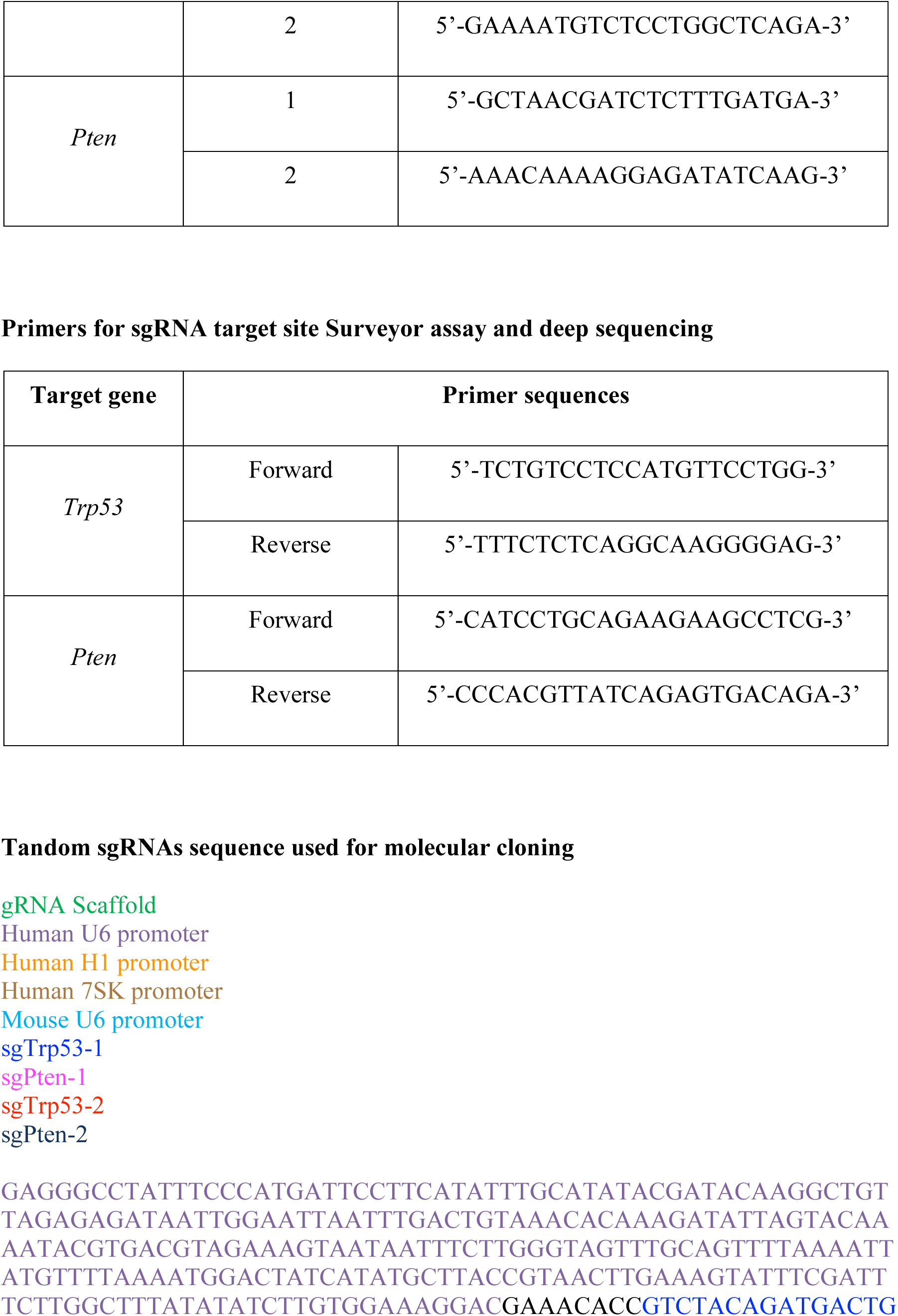
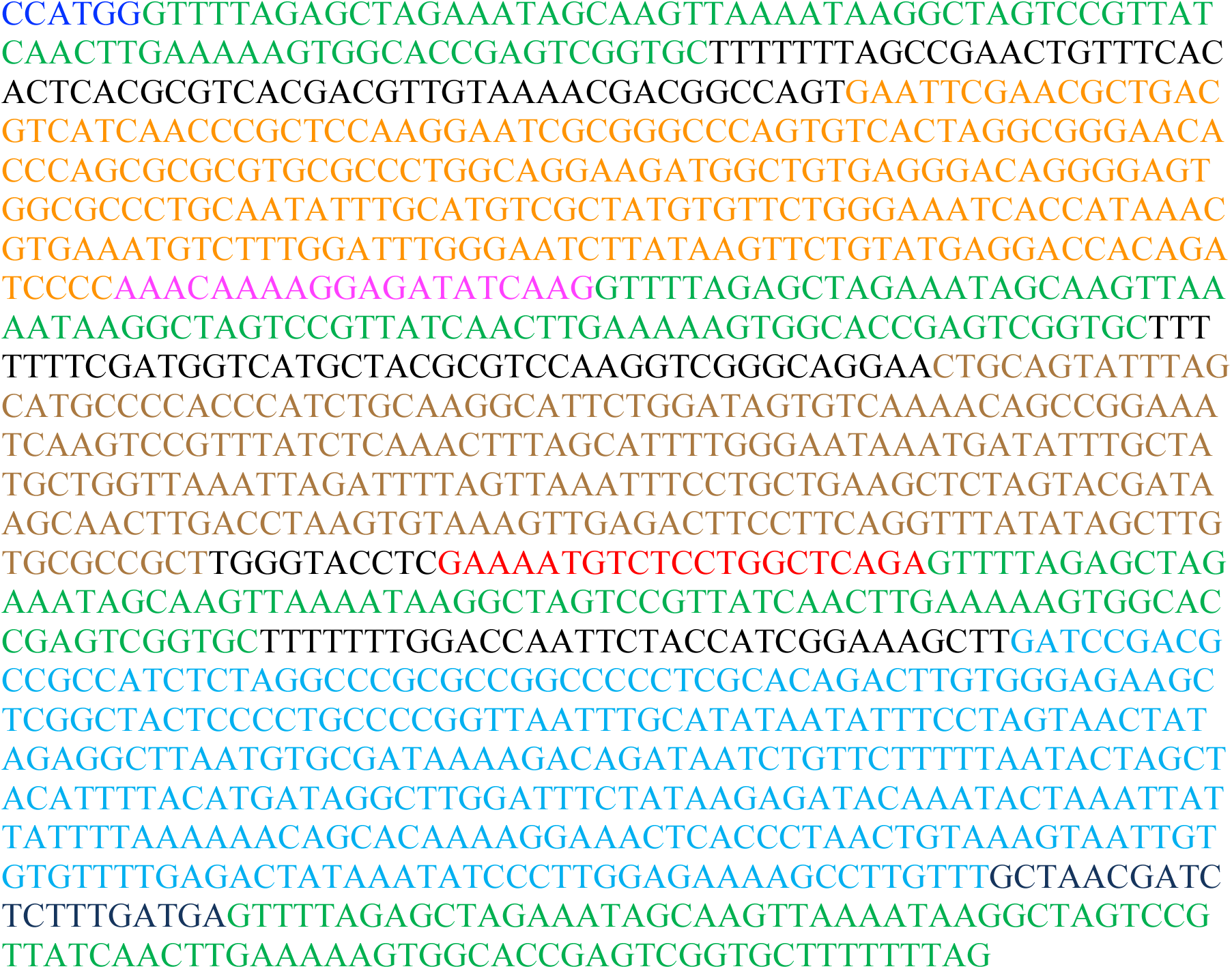
Primers and sgRNA sequences in this study Primers for mouse genotyping.

**Table S2.** List of *Bas-trace* and *Lum-trace* mice tumor phenotypes. ‘*’ represents the number of transformed lesions (N): N >=10 ‘***’; 5<N<10 ‘**’; N<=5 ‘*’ ‘P’ represents the number of protruding tumor clones into the lumen, which have more than 5 layers of tumor cells.

| *Bas-trace* |  |  |  |
| --- | --- | --- | --- |
| Mouse ID (#) | 3 months post-electroporation |  |  |
|  | Precancerous | Type I | Type II |
| #1087 | *** | **<br>P=0 | *<br>P=0 |
| #1092 | *** | **<br>P=0 | **<br>P=0 |
| #1093 | ** | *<br>P=0 | *<br>P=0 |
| #1128 | *** | ***<br>P=4 | ***<br>P=0 |
| #1129 | *** | **<br>P=2 | **<br>P=0 |
| #7811 | *** | ***<br>P=4 | ***<br>P=0 |
| #7815 | * | *<br>P=2 | *<br>P=0 |
| #7845 | *** | ***<br>P=2 | ***<br>P=0 |
| #7849 | *** | ***<br>P=0 | ***<br>P=0 |
| Mouse ID (#) | 6 months post-electroporation |  |  |
|  | Precancerous | Type I | Type II |
| #1019 | *** | ***<br>P=5 | ***<br>P=1 |
| #1028 | *** | **<br>P=3 | ***<br>P=1 |
| #1891 | *** | **<br>P=2 | ***<br>P=1 |
| #1892 | *** | *<br>P=0 | ***<br>P=0 |
| #1917 | *** | **<br>P=6 | ***<br>P=4 |
| #1922 | ** | ***<br>P=12 | ***<br>P=4 |
| #1923 | *** | ***<br>P=2 | ***<br>P=0 |
| #1984 | *** | **<br>P=4 | **<br>P=0 |
| #1988 | *** | **<br>P=2 | ***<br>P=0 |
| #9870 | * | **<br>P=4 | **<br>P=3 |
| #9875 | *** | ***<br>P=2 | ***<br>P=2 |

| *Lum-trace* |  |  |  |
| --- | --- | --- | --- |
| Mouse ID (#) | 3 months post-electroporation |  |  |
|  | Precancerous | Type I | Type II |
| #1174 | *** | **<br>P=2 | ***<br>P=0 |
| #1176 | *** | **<br>P=0 | ***<br>P=0 |
| #1482 | *** | **<br>P=3 | **<br>P=0 |
| #1836 | *** | ***<br>P=2 | ***<br>P=0 |
| #1838 | *** | ***<br>P=8 | ***<br>P=0 |
| #7957 | *** | ***<br>P=7 | ***<br>P=0 |
| #9396 | *** | **<br>P=5 | ***<br>P=0 |
| Mouse ID (#) | 6 months post-electroporation |  |  |
|  | Precancerous | Type I | Type II |
| #1012 | *** | **<br>P=3 | ***<br>P=2 |
| #1723 | *** | ***<br>P=12 | ***<br>P=1 |
| #1724 | *** | ***<br>P=23 | ***<br>P=3 |
| #1725 | *** | ***<br>P=9 | ***<br>P=5 |
| #9230 | *** | ***<br>P=8 | ***<br>P=8 |
| #9558 | ** | *<br>P=3 | ***<br>P=4 |
| #9758 | * | ***<br>P=15 | ***<br>P=2 |
| #9759 | *** | **<br>P=1 | ***<br>P=0 |
| #9762 | * | ***<br>P=19 | ***<br>P=12 |

**Table S3.** Primary antibodies used for IF and IHC staining.

| **Antigen** | **Supplier** | **Ig type** | **Dilution** |
| --- | --- | --- | --- |
| CK14 | Biolegend #906004 | Chicken IgY | 1:1000 |
| Ki67 | DakoCytomation #M7249 | Rat IgG2a | 1:500 |
| tdTomato | OriGene #AB8181-200 | Goat IgG | 1:1000 |
| CK5 | Covance #PRB-160P | Rabbit IgG | 1:1000 |
| CK18 | Abcam #ab668 | Mouse IgG1 | 1:200 |
| FoxA1 | Abcam #ab23738 | Rabbit IgG | 1:100 |
| GFP | Abcam #A13970 | Chicken IgY | 1:4000 |
| Gata3 | eBioscience #14-9966-80 | Rat IgG2b | 1:100 |
| Pten | Cell Signalling #9559T | Rabbit IgG | 1:100 (IHC) |
| pAKT | Cell Signalling #3787S | Rabbit IgG | 1:100 (IHC) |
| Trp53 | Abcam #ab241566 | Rat IgG2a | 1:100 |

